# Non-disruptive inducible labeling of ER-membrane contact sites using the Lamin B Receptor

**DOI:** 10.1101/2024.05.31.596797

**Authors:** Laura Downie, Nuria Ferrandiz, Megan Jones, Stephen J. Royle

## Abstract

Membrane contact sites (MCSs) are areas of close proximity between organelles that allow the exchange of material, among other roles. The endoplasmic reticulum (ER) has MCSs with a variety of organelles in the cell. MCSs are dynamic, responding to changes in cell state, and are therefore best visualized through inducible labeling methods. However, existing methods typically distort ER-MCSs, by expanding contacts or creating artificial ones. Here we describe a new method for inducible labeling of ER-MCSs using the Lamin B receptor (LBR) and a generic anchor protein on the partner organelle. Termed *LaBeRling*, this versatile, one-to-many approach allows labeling of different types of ER-MCSs (mitochondria, plasma membrane, lysosomes, early endosomes, lipid droplets and Golgi), on-demand, in interphase or mitotic cells. LaBeRling is non-disruptive and does not change ER-MCSs in terms of the contact number, extent or distance measured; as determined by light microscopy or a deep-learning volume electron microscopy approach. We applied this method to study the changes in ER-MCSs during mitosis and to label novel ER-Golgi contact sites at different mitotic stages in live cells.

## Introduction

In eukaryotic cells, membrane contact sites (MCSs) are areas of close proximity between two membranes of different identity. MCSs allow the exchange of material between the two membranes without fusion, and they also regulate organelle positioning, dynamics, and number (Scorrano et al., 2019). The ER occupies a large volume of the cell and forms contacts with multiple membranes of different identity (Wu et al., 2018). ER-MCSs therefore play a central role in cellular communication across organelles; they are also dynamic and must adapt in response to changes in cell state. The ER and other membrane compartments remodel upon entry into mitosis (Carlton et al., 2020). For example, the nuclear envelope breaks down and the Golgi fragments (Ungricht and Kutay, 2017; Shorter and Warren, 2002). How ER-MCSs change in response to this remodeling is not well understood. Recent work indicates that changes in ER-MCSs may be coordinated with other mitotic processes through regulation of tethering proteins in mitosis (James et al., 2024). The study of MCSs is hampered by the lack of live cell labeling methods that i) specifically label MCSs, ii) do not interfere with contacts, and iii) allow visualization of contacts during specific cell cycle stages.

ER-MCSs were first observed in electron microscopy (EM) studies in fixed cells (Bernhard and Rouiller, 1956; Copeland and Dalton, 1959; Lewis and Tata, 1973; Henkart et al., 1976; Scorrano et al., 2019). Other visualization methods include those based on measures of organelle membrane proximity from live cell multispectral imaging (Valm et al., 2017) or fixed super-resolution 3D images (Cardoen et al., 2024). Ideally, MCS labeling methods in live cells must distinguish MCS from regions where membranes are in close proximity by chance, as recently reviewed by Nakatsu and Tsukiji (2023). Examples for visualizing ER-MCSs include proximity-dependent methods, where fluorescence is produced when protein tags on each membrane are in close proximity: fluorescence resonance energy transfer (FRET) (Venditti et al., 2019), split fluorescent proteins (FPs) (Yang et al., 2018; Calì and Brini, 2021), and dimerization-dependent FPs (ddFPs) (Miner et al., 2024). Synthetic constructs based on tethering proteins have also been used to visualize ER-MCSs (Chang et al., 2013). A major limitation of these methods is that labeling is not inducible. To study MCSs during mitosis, labeling that persists and potentially stabilizes interphase contacts is undesirable.

Inducible methods allow temporal control of MCS labeling, and typically use tagged proteins on each apposing membrane that heterodimerize in response to light or a chemical (Sittewelle et al., 2023; Nakatsu and Tsukiji, 2023; Casas et al., 2023; Li et al., 2024). However, the initial expression and/or the induced heterodimerization of these proteins usually increase the contact between membranes, through expanding the area of preexisting MCSs or by creating artificial tethers between the ER and the apposing membrane. Expansion of ER-mitochondria MCSs over time after the inducing labeling has been observed in live cells and by EM (Csordás et al., 2010; Komatsu et al., 2010). Here the ER could be seen wrapping around the mitochondria surface; increasing ER-mitochondria coverage from ∼10 % to ∼30–90 % after induction (Csordás et al., 2010). Moreover, an optogenetic approach also showed an increase of ∼20 % in ER/mitochondria signal overlap in live HeLa cells; and a larger increase in ER-lysosome contacts using this approach in live Cos-7 cells (Benedetti et al., 2020). ER-plasma membrane (PM) contacts were also increased after labeling using rapamycin-induced heterodimerization (Várnai et al., 2007). In fact, we previously exploited this property to artificially “glue” the ER to the PM during mitosis in order to free chromosomes trapped by the ER (Ferrandiz et al., 2022). This limits the use of inducible methods for the study of MCSs, as labeling manipulates the ER-MCSs.

Our aim was to develop an inducible labeling system which allows fast and specific labeling of ER-membrane contacts, without disrupting the contacts themselves. Using chemical-induced heterodimerization with a tagged anchor protein at the target membrane, we serendipitously found that Lamin B receptor (LBR), which localizes to the ER and inner nuclear envelope, specifically labeled ER-MCSs upon relocalization. We called this method *LaBeRling*. We found that LaBeRling caused no detectable change in the contact number, extent or distance measured from high resolution 3D EM datasets. It can be used to label multiple different ER-MCS types and we applied this method to study MCSs in mitosis and use it to reveal novel ER-Golgi MCSs in live mitotic cells.

## Results

### LBR forms clusters after relocalization to the plasma membrane

In principle, induced heterodimerization of proteins tagged with FKBP and FRB domains can be used to label membrane contact sites on-demand. The advantage to this method is that the moment of labeling is controlled by the investigator, a major disadvantage is that the heterodimerization may distort existing contact sites or even induce new, or artificial contacts (Figure 1A). We began by comparing the relocalization of two different proteins: Sec61β and the lamin B receptor, LBR; to the plasma membrane (PM) using induced heterodimerization. Both Sec61β and LBR are found in the ER, with LBR being additionally present on the inner nuclear envelope in interphase. Using Stargazin-dCherry-FRB as a plasma membrane anchor, we found that FKBP-GFP-Sec61β relocalization causes the ER to become glued to the plasma membrane in interphase or during mitosis, as reported previously (Ferrandiz et al., 2022). By contrast, LBR-FKBP-GFP was found in discrete clusters at the plasma membrane following rapamycin addition (Figure 1B) The formation of LBR-FKBP-GFP clusters was dependent on both the expression of the plasma membrane anchor and on the addition of rapamycin (Supplementary Figure S1).

**Figure 1.**
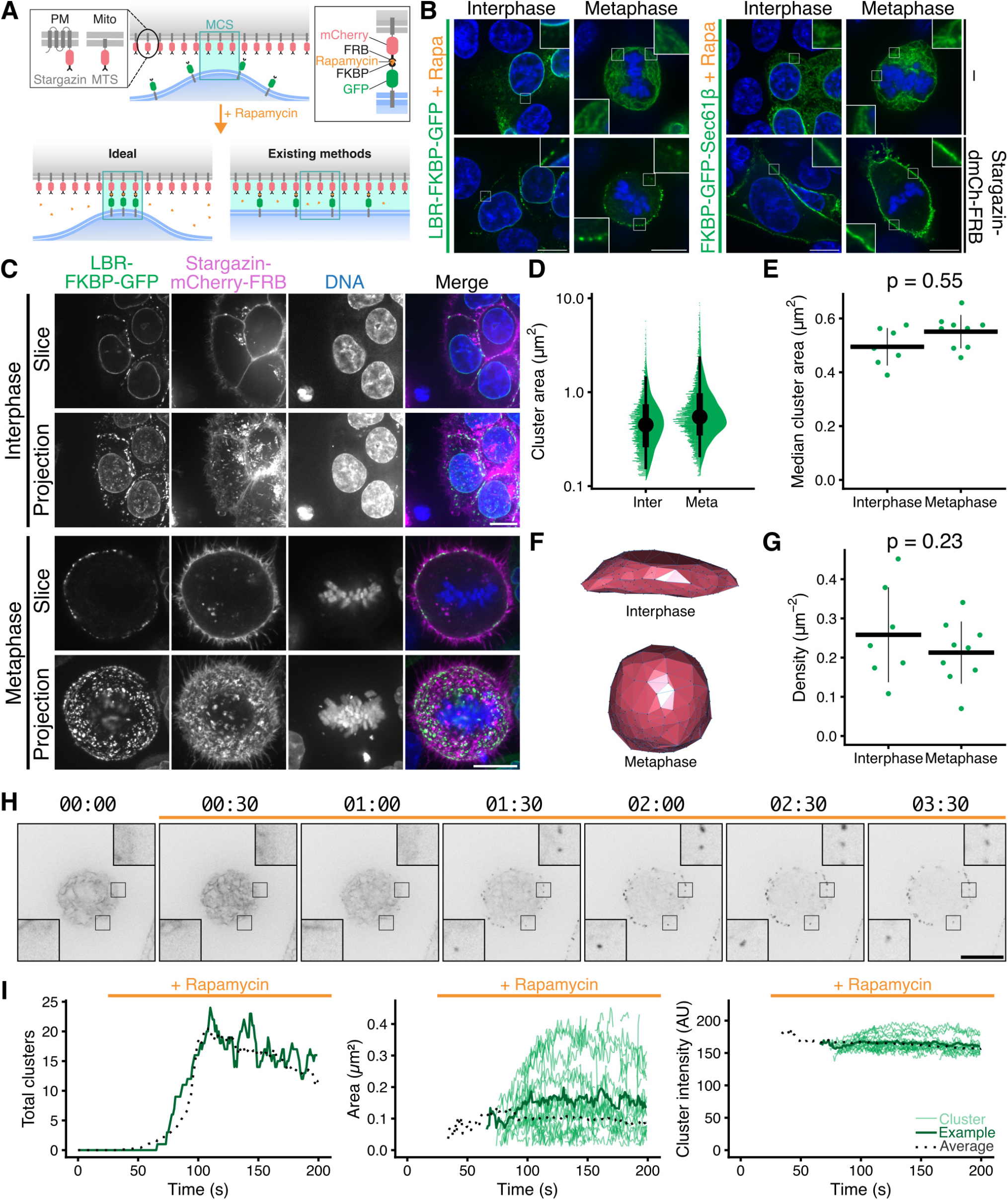
Properties of LBR-FKBP-GFP clusters at the plasma membrane following induced relocalization. (**A**) Schematic diagram to show heterodimerization of FKBP and FRB domains with rapamycin and how this may be used to label ER-membrane contact sites, but also how this method may distort contact sites. Membrane contact site (MCS) is labeled by a blue box and a green shading is used indicate membranes within close distance of each other. (**B**) Example micrographs of HCT116 cells co-expressing LBR-FKBP-GFP or FKBP-GFP-Sec61β (green) and optionally Stargazin-dCherryFRB, each treated with rapamycin (200 nM) for 30 min before fixation and DNA staining (blue). Scale bars, 10 μm; Insets, 3× expansion of ROI. Note, additional controls are shown in full in Supplementary Figure S1A. (**C**) Micrographs of typical HCT116 LBR-FKBP-GFP (green) knock-in cells expressing Stargazin-mCherry-FRB (magenta), stained with SiR-DNA (blue), treated with rapamycin (200 nM). Single confocal slices or z-projections for cells in interphase or mitosis are shown. (**D**) Raincloud plot to show size distribution of LBR-FKBP-GFP clusters analyzed in 3D. (**E**) Plot of the median contact area for each cell (dots). Mean and ± sd are indicated by crossbar. (**F**) 3D cell surface approximation generated using the location of all segmented LBR-FKBP-GFP clusters (see Methods). (**G**) Plot of the density of clusters (total clusters divided by cell surface). Dots show cells, mean and ± sd are indicated by crossbar. (**H**) Stills from a movie of LBR-FKBP-GFP cluster formation upon rapamycin addition (200 nM, orange bar). Cells were as described in C. Timescale, mm:ss. Scale bars, 10 μm; Insets, 3× expansion of ROI. (**I**) Plots to show the number, size, and intensity of LBR-FKBP-GFP clusters in a single slice over time. Thin green lines and dark green line, clusters from and average for the cell shown in F; black dotted line, average of 14 different cells.

To avoid the possibility that overexpression of LBR-FKBP-GFP contributed to cluster formation, we generated a knock-in cell line where LBR was tagged with FKBP-GFP at its endogenous locus (Supplementary Figure S2). Using these cells, we confirmed that the formation of LBR-FKBP-GFP clusters occurred in a similar way (Supplementary Figure S1). Furthermore, similar LBR-FKBP-GFP clusters were seen when using SH4-FRB-EBFP2, a peripheral membrane protein, in place of the multipass Stargazin plasma membrane anchor (Supplementary Figure S3), which suggested that the identity of the anchor was unimportant and that the difference in behavior could be attributed solely to the ER-resident protein. Interestingly, there was minimal clustering of the plasma membrane anchor upon relocalization of LBR-FKBP-GFP (Figure S3A and Figure 1C). We also found no evidence for co-clustering of mCherry-Sec61β or of LBR-mCherry when LBR-FKBP-GFP was relocalized to the plasma membrane, which suggested that the clusters did not represent non-specific aggregates of ER or LBR protein itself (Figure S3). Visualization of the ER using PhenoVue Fluor568-ConA revealed no gross changes in ER morphology when clustering of LBR-FKBP-GFP was induced (Supplementary Figure S4). Indeed because the ER stayed intact while LBR-FKBP-GFP clustered in the ER at discrete sites on the plasma membrane, it suggested that this manipulation may be labeling ER-PM contact sites.

### Properties of LBR-FKBP-GFP clusters at the plasma membrane following relocalization

We next investigated the properties of the clusters that form after inducing the relocalization of LBR-FKBP-GFP to the plasma membrane. We characterized the properties of the LBR-FKBP-GFP clusters in cells in interphase or at metaphase using 3D segmentation of confocal z-stacks of HCT116 LBR-FKBP-GFP knock-in cells expressing Stargazin-mCherry-FRB treated with rapamycin (200 nM, 20 min) (Figure 1C). The surface area of the clusters (see Methods for definition) was variable but the median cluster area per cell was (0.50 ± 0.07, interphase; 0.55 ± 0.06 μm^2^, metaphase), with no significant difference between interphase and metaphase cells (Figure 1D,E). Hundreds of clusters were detected in each cell, but to normalize for differences in cell size and understand the density of LBR-FKBP-GFP clusters at the plasma membrane a cell surface approximation was generated and the density of clusters per unit area was determined (Figure 1F,G). Again the density of clusters was similar in interphase and metaphase cells (0.26 ± 0.12, interphase; 0.21 ± 0.08 μm^−2^, metaphase). Given the average size of the clusters and their density, the coverage of the plasma membrane was ∼10 %, which is similar to published estimates of PM coverage with ER-PM contact sites (Wu et al., 2017).

To investigate the formation and dynamics of LBR-FKBP-GFP clusters, we imaged mitotic cells during the induced relocalization of LBR-FKBP-GFP to the plasma membrane (Figure 1H). These movies revealed that multiple clusters that were distributed around the cell, formed simultaneously and with similar kinetics (Supplementary Video SV1). The clusters began to form ∼90 s after rapamycin addition, with typically less than 20 clusters in a single confocal slice, and persisted throughout the duration of the movie (up to 5 min). The clusters that formed did so at the expense of fluorescence in the ER which “drained away” with similar kinetics.

Using a spot-tracking procedure we could monitor the behavior of each cluster. Analysis revealed that once formed, the total number, size, and brightness of the clusters were essentially constant on this time scale (Figure 1I). There were very few splitting or merging of clusters, and any appearance or disappearance of clusters could be attributed to movement into or out of the imaging plane. Overall, the mobility of the clusters was very low (0.016 μm s^−1^, median, n = 16 cells).

The coordinated appearance and stability of fluorescence suggests that relocalized LBR-FKBP-GFP labels pre-existing ER-PM contact sites.

### Relocalized LBR-FKBP-GFP labels pre-existing ER-PM contact sites

Are the LBR-FKBP-GFP clusters that form after relocalization, coincident with pre-existing ER-PM contact sites? To answer this question we took confocal z-stacks of interphase and mitotic cells before and after LBR-FKBP-GFP relocalization to Stargazin-EBFP2-FRB using rapamycin (200 nM). To identify the ER-PM contact sites, we used a modified MAPPER construct (mScarlet-I3-6DG5-MAPPER) which is an established marker of ER-PM contact sites (Chang et al., 2013). From these images, we could clearly see colocalization between the contact sites marked by MAPPER and the clusters of relocalized LBR-FKBP-GFP in interphase and mitotic cells (Figure 2A). This was also true of the relocalization captured in live HCT116 or HeLa cells (Supplementary Video SV2, SV3). The contact sites where colocalization occurred were present before LBR-FKBP-GFP was relocalized, indicating that labeling is of pre-existing contact sites.

**Figure 2.**
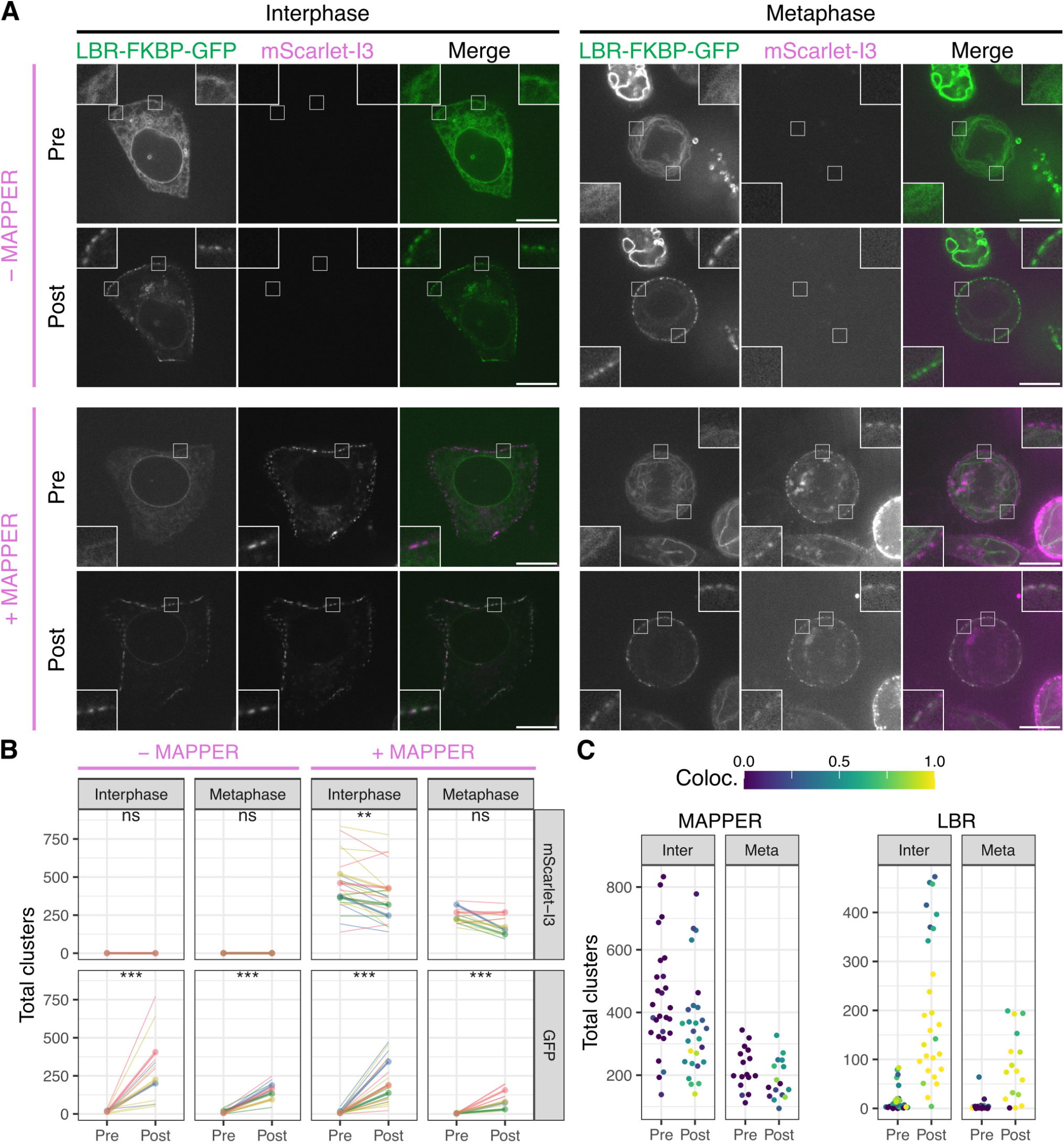
Relocalized LBR-FKBP-GFP labels pre-existing ER-PM contact sites. (**A**) Example micrographs of HCT116 cells co-expressing LBR-FKBP-GFP (green) and Stargazin-EBFP2-FRB (not shown) and optionally MAPPER (mScarlet-I3-6DG5-MAPPER, magenta) as indicated. A single slice of a z-stack of the same cell is shown before (pre) or after (post) rapamycin (200 nM, 20 min) addition. Scale bars, 10 μm; Insets, 3× expansion of ROI. (**B**) Comparison of total 3D clusters per cell detected in mScarlet-I3 (MAPPER) or GFP (LBR-FKBP-GFP) channels. Thin lines indicate the pre and post values for each cell. Thick lines and dots indicate the average per experimental repeat. Color indicates experimental repeat. Paired t-tests with Holm-Bonferroni correction for multiple testing: ns, not significant; ***, p < 0.001; **, p < 0.01. (**C**) Colocalization analysis. For cells co-expressing MAPPER, the number of MAPPER or LBR clusters is shown and the extent that these clusters colocalize with LBR or MAPPER clusters is indicated by the colorscale.

3D segmentation and quantification of the contact sites marked by MAPPER and the clusters of LBR-FKBP-GFP confirmed that the formation of LBR-FKBP-GFP clusters by inducing relocalization was significant. There was no change in the number of MAPPER clusters per cell in mitotic cells, and a small but significant decrease in interphase which was probably attributable to photobleaching (Figure 2B). Importantly, we found no evidence for increases in MAPPER clusters after LBR-FKBP-GFP relocalization which would have indicated the artificial formation of new contact sites. When the colocalization of the two fluorescence channels was measured we found in most cells LBR-FKBP-GFP clusters were contact sites (Figure 2C). The fraction of contact sites that were labeled by LBR-FKBP-GFP was lower, a result which may have been influenced by photobleaching of GFP vs mScarlet-I3.

We found no influence of MAPPER expression on the number of LBR-FKBP-GFP clusters that form upon relocalization to the plasma membrane. For example, there were 213.4 ± 89.4 LBR-FKBP-GFP clusters in MAPPER expressing interphase cells compared with 277.2 ± 112.6 in those not co-expressing MAPPER (p = 0.7, Tukey’s post-hoc test). This indicates that there is no interference between the two labeling types.

Finally, this dataset also revealed that there are fewer ER-PM contact sites at metaphase than there are in interphase (Figure 2B). for example, the number of MAPPER clusters in interphase and metaphase cells, before rapamycin treatment was significantly lower (429.3 ± 74.5, interphase; 258.5 ± 45.3, metaphase; p = 0.002). A pattern repeated for post-rapamycin treatment (p = 0.002) and for LBR-FKBP-GFP clusters in MAPPER expressing cells post-rapamycin (p = 0.04). In summary, we could confirm that LBR-FKBP-GFP relocalization to the plasma membrane labels pre-existing ER-PM contact sites and that there were no additional sites created by this relocalization. We also documented a decrease in the number of contacts in mitotic cells compared with non-dividing cells.

### Using relocalization of LBR-FKBP-GFP to detect functional changes in ER-PM membrane contact sites

Application of the ER calcium pump inhibitor, thapsigargin, was reported to increase the density of ER-PM MCSs (Chang et al., 2013). To test if the relocalization of LBR-FKBP-GFP to the plasma membrane was sensitive to functional changes in membrane contact sites, we applied thapsigargin (1 μM) for 20 min prior to application of rapamycin. Measurement of the resulting LBR-FKBP-GFP clusters showed a significant increase in coverage at the plasma membrane (Supplementary Figure S5). These experiments suggest that the relocalization of LBR-FKBP-GFP to the plasma membrane can be used to report changes in MCS size when ER calcium signaling is manipulated.

### Relocalization of LBR-FKBP-GFP to mitochondria highlights ER-mitochondria contact sites

Having established that LBR can be used to inducibly label ER-PM contact sites, we next tested if it could be used to label ER-mitochondria contact sites. HCT116 cells transiently co-expressing MitoTrap (Mito-mCherry-FRB) and either LBR-FKBP-GFP or FKBP-GFP-Sec61β were imaged in interphase or mitosis (Figure 3A). Following relocalization with rapamycin (200 nM), LBR-FKBP-GFP was clustered at discrete sites on each mitochondrion with typically one contact per mitochondrion visible by light microscopy. By contrast, FKBP-GFP-Sec61β fluorescence completely surrounded each mitochondrion suggesting that the relocalization of this protein caused the ER to wrap around the mitochondria (Figure 3A). This pattern was similar in interphase or mitotic cells, and mirrored the observations previously using a plasma membrane anchor.

**Figure 3.**
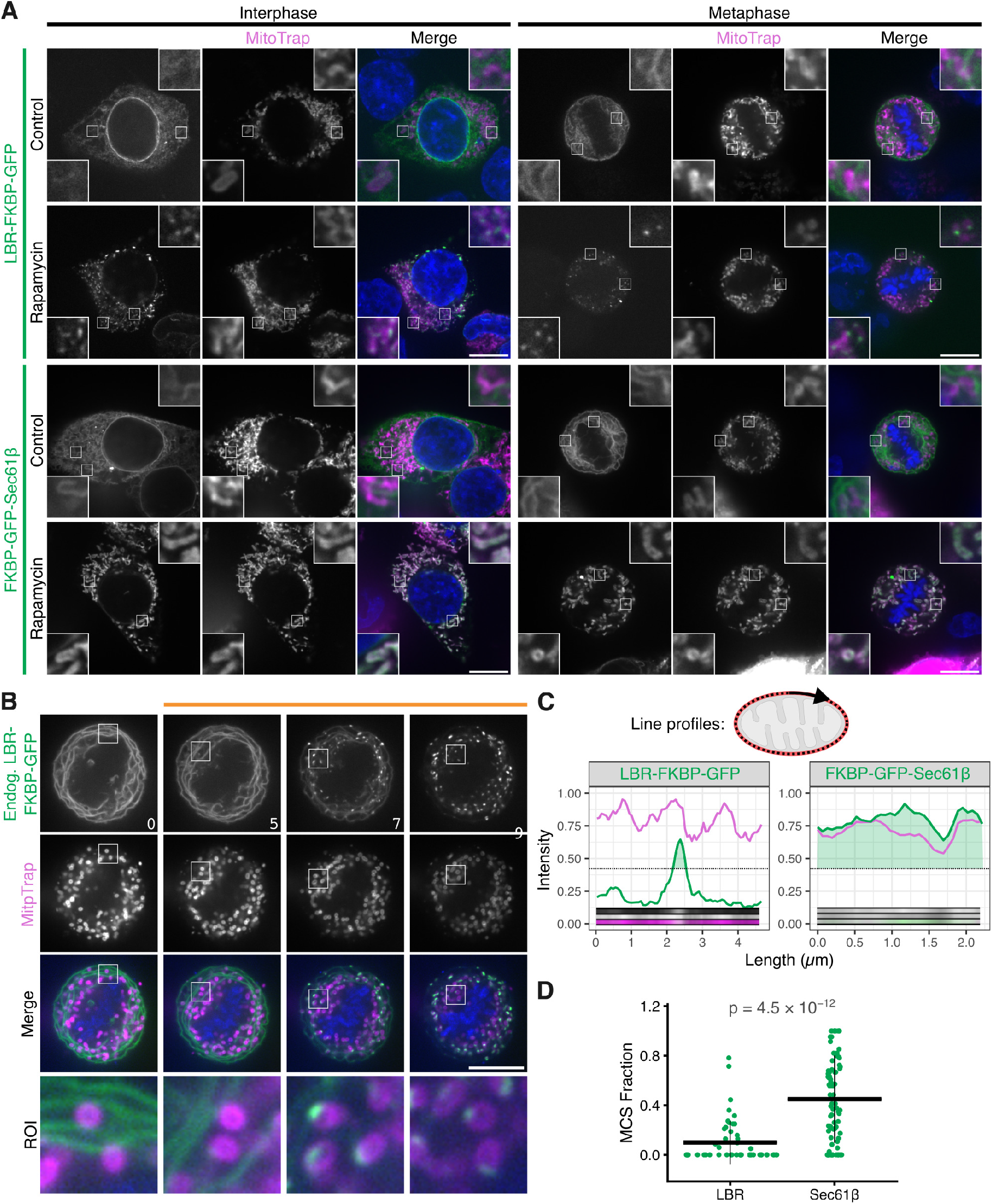
Relocalization of LBR-FKBP-GFP to mitochondria highlights ER-mitochondria contact sites. (**A**) Example micrographs of HCT116 cells expressing either LBR-FKBP-GFP or FKBP-GFP-Sec61β (green) and MitoTrap (Mito-mCherry-FRB, magenta). Relocalized samples were treated with rapamycin (200 nM) for 30 min before fixation. Control samples were not treated with rapamycin. A single slice from a z-stack of an interphase or metaphase cell are shown. Scale bars, 10 μm; Insets, 4× expansion of ROI. (**B**) Stills from a live cell imaging experiment with HCT116 LBR-FKBP-GFP (green) knock-in cells co-expressing MitoTrap (magenta), treated with rapamycin (200 nM) as indicated. Time, min; Scale bars, 10 μm; Zooms, 6.5× expansion of ROI. (**C**) Line profiles measured around the mitochondrial perimeter from mitotic cells in A, as represented in the schematic. Plots show the intensity of LBR-FKBP-GFP or FKBP-GFP-Sec61bβ (green) and MitoTrap (magenta) signal measured around an individual mitochondria. Dotted line indicates the threshold for segmentation, light green area indicates sections of the line above threshold. Insets show the line profile images. (**D**) Plot of the fraction of the mitochondrial perimeter that is above threshold. Quantification is of line profiles like those shown in C. P-value, Tukey’s HSD *post hoc* test; n_cell_ = 3 (LBR), 8 (Sec61β).

The clusters of LBR-FKBP-GFP on mitochondria following relocalization were also observed with the endogenously tagged protein in live cells, again suggesting that the cluster formation was not an artifact of overexpression or fixation (Figure 3B). Live-cell imaging of the relocalization revealed that cluster formation was rapid (∼20 s) and came at the expense of fluorescence in the ER (Supplementary Video SV4). Wrapping of mitochondria by FKBP-GFP-Sec61β occurred on longer timescale (Supplementary Video SV5). We used fluorescence profiles to determine the fraction of the organelle that was occupied by GFP signal (Figure 3C), this analysis revealed significantly higher coverage by FKBP-GFP-Sec61β compared to LBR-FKBP-GFP (Figure 3D).

The appearance of discrete clusters of LBR-FKBP-GFP upon relocalization to mitochondria suggested that these clusters most likely represent ER-mitochondria contact sites.

### Inducible labeling of ER-mitochondria contacts using LBR does not affect ER-mitochondria contacts

Are ER-mitochondria contacts altered by the relocalization of LBR-FKBP-GFP to MitoTrap? As there was no obvious choice of marker to assess this, we used a 3D-EM approach to examine all ER-mitochondria contacts (Figure 4). We first imaged live HCT116 LBR-FKBP-GFP knock-in cells co-expressing MitoTrap and confirmed the relocalization of LBR-FKBP-GFP to mitochondria or not in the case of the control (no rapamycin, Figure 4A). The same cell was then processed for SBF-SEM and relocated for imaging by 3D-EM. We used a machine learning approach to infer in 3D the mitochondria and ER in all of the resulting datasets (control, 3; rapamycin, 4) using a manually segmented subvolume from one dataset (Figure 4B). Next, using the inferred maps of mitochondria and ER, we used an automated procedure to detect the regions of ER that were within a defined distance of the mitochondria, and then segment those to measure the size of the ER region that is in contact with the mitochondrion (see Methods). We verified that the procedure identified genuine ER-mitochondrion contacts by sampling a number of contacts and by visualizing them in 3D (Figure 4B). Using the resulting data we could measure the total ER contacts per mitochondrion and plot the total contact area as a function of mitochondrion surface area (Figure 4C). At all defined distances assessed, from 10–50 nm, we found that the contacts were similar between the two experimental groups (Figure 4D). These data indicate that relocalization of LBR-FKBP-GFP can be used to discretely label ER-mitochondria contact sites, without affecting the morphology of those contacts. Since LBR labeling of ER-PM contacts was shown to be similarly non-invasive, we propose that our method is an innocuous way to label contact sites inducibly. We call this method, LaBeRling.

**Figure 4.**
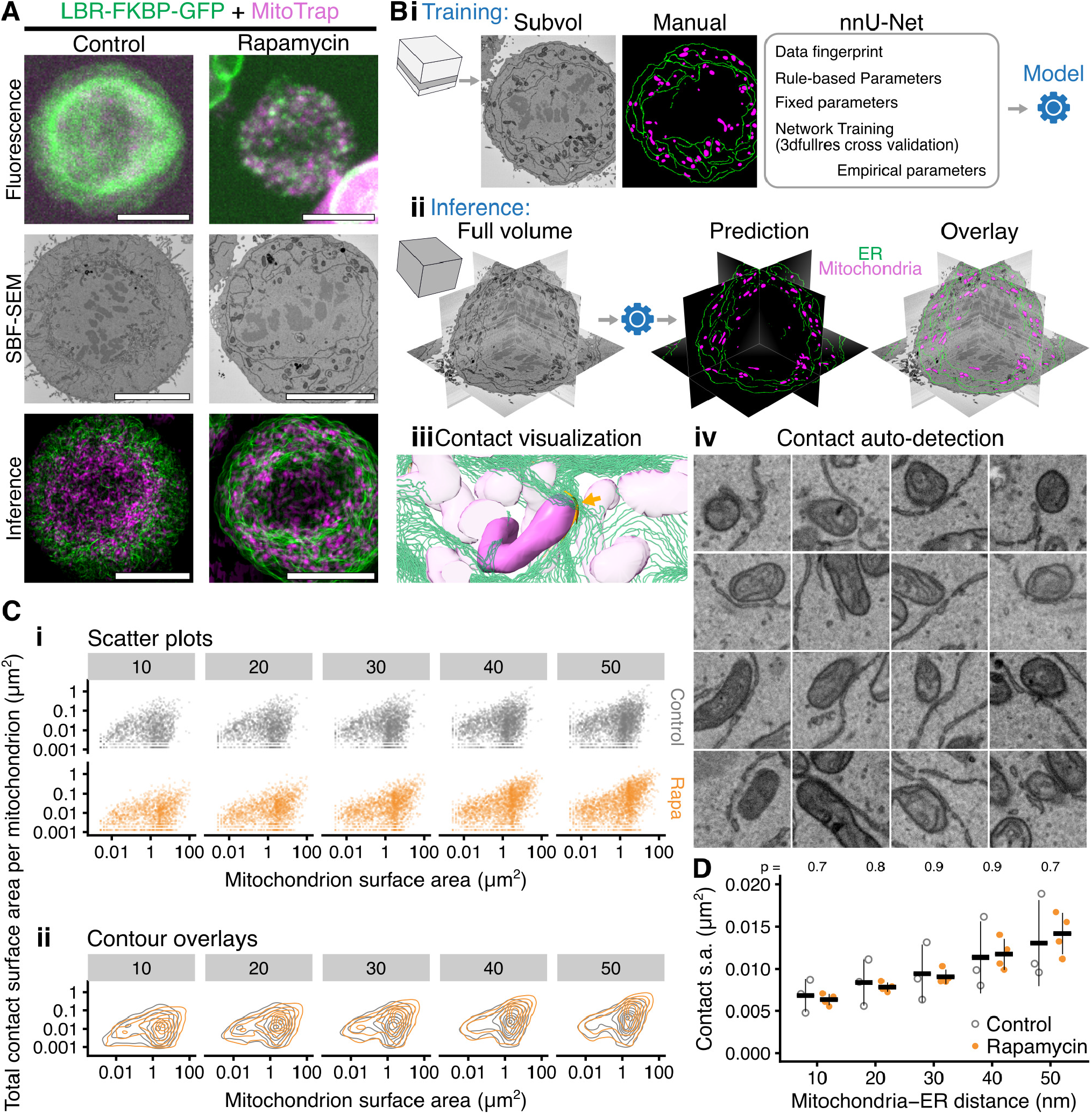
Inducible labeling of ER-mitochondria contacts using LBR does not affect ER-mitochondria contacts. (**A**) Example micrographs of HCT116 LBR-FKBP-GFP (green) knock-in cells expressing MitoTrap (magenta) treated with Rapamycin (200 nM, 30 min) or not (Control). Mitotic stage was confirmed by imaging SiR-DNA (not shown). Following fixation and processing, the same cell was imaged again by serial block face-scanning electron microscopy (SBF-SEM). Finally, SBF-SEM datasets were used to infer the location of ER (green) and mitochondria (magenta), a Z-projection is shown. Scale bar, 10 μm. (**B**) Large-scale machine learning segmentation of ER and mitochondria from SBF-SEM data. (i) Supervised training of nnU-Net using a subvolume of one SBF-SEM dataset in 3dfullres mode. The resulting model is then used to infer the location of ER and mitochondria in the whole volume of multiple datasets. (ii) An image analysis pipeline (see Methods) detects the ER-mitochondria contact areas that are equal or less than the search distance (10–50 nm) from the nearest mitochondrion. (iii) The contacts may be visualized in 3D: orange contact is shown at a highlighted mitochondrion (arrow), ER is represented by green contour lines for clarity. (iv) Contacts detected can be mapped back to the original data for verification. A random selection of contacts from the 50 nm search distance collection are shown. (**C**) Plots of the total ER-mitochondria contact surface area per mitochondrion vs the surface area of the mitochondrion, for each search distance (indicated in gray box, nm). (i) scatter plots, n_mito_ = 1494-2746 (control), 1432-2421 (rapamycin); 10-50 nm. (ii) contour plots are shown overlaid to compare the distribution between control and rapamycin. (**D**) Mean ER-mitochondria contact surface area per cell for each search distance. Each cell is represented as a dot, the mean ± sd is shown by a crossbar; n_cell_ = 3 (control), 4 (rapamycin); p values from Student’s t-test with Welch’s correction.

### General application of the LaBeRling method

Having demonstrated LaBeRling in HCT116 cells, we wanted to test if the method worked in other cell types. We tested LaBeRling of ER-PM and ER-mitochondria MCSs in three cell lines: RPE-1, Cos-7 and HeLa. In each case relocalization of LBR-FKBP-GFP to its respective anchor using rapamycin (200 nM) induced discrete puncta formation in each cell type (Supplementary Figure S6). This suggests that the method may be broadly applicable in a variety of cell lines.

So far we have used up to 30 min rapamycin addition to induced LaBeRling. For most experiments, this will be sufficient, however for long-term highlighting of membrane contact sites, the use of rapamycin as an induction agent may be problematic due to its bioactivity on the timescale of hours. Accordingly, we tested whether rapalog AP20187 could similarly be used to induce LaBeRling. Using coexpression of either Stargazin-mCherry-FRB(T2098L) or Mito-mCherry-FRB(T2098L) variants with LBR-FKBP-GFP, we saw similar highlighting of ER-PM or ER-mitochondria MCSs, respectively (Supplementary Figure S7).

We next attempted long-term LaBeRling of ER-PM MCSs in HeLa cells. Following induction, we could see that contacts remained brightly fluorescent up to 4 h (Supplementry Figure S8). Therefore we tested whether such long-term LaBeRling was detrimental to cell health. Using this method we could follow cells with LaBeRled ER-PM MCSs for up to ∼9 h. Cells migrated similarly to their respective controls and the MCSs remained fluorescent throughout (Supplementary Video SV6 and SV7). These observations suggest that the method can be used in other contexts beyond shortterm highlighting of contact sites, and that LaBeRling is effective in other cell lines.

### Relocalized LBR-FKBP-GFP can mark several different ER-membrane contact types

As LaBeRling can be used to highlight ER-PM and ER-mitochondria contact sites, we next asked if it could be applied to similarly label other ER-membrane contact sites. To do this we tested three additional anchor proteins: perilipin-3 (PLIN3), early endosome antigen 1 (EEA1), and lysosome-associated membrane glycoprotein 1 (LAMP1); that mark lipid droplets, early endosomes, and lysosomes, respectively. We compared the relocalization of LBR-FKBP-GFP with that of FKBP-GFP-Sec61β to differentiate genuine contact site labeling from non-specific recruitment of ER to the target membrane.

At early endosomes and lysosomes, LBR-FKBP-GFP was relocalized to clusters whereas the relocalization of FKBP-GFP-Sec61β matched the fluorescence of the anchor (Figure 5A). At lipid droplets however, relocalization of either LBR-FKBP-GFP or FKBP-GFP-Sec61β caused a coincidence of the GFP fluorescence with FRB-mCherry-PLIN3 that was not distinguishable between the two ER proteins (Figure 5A). Analysis of the MCS fraction from organelle fluorescence profiles supported these observations (Figure 5B and Supplementary Figure S9). This suggested that LaBeRling of discrete ER-endosome or ER-lysosome contacts was possible, while Sec61β relocalization engulfed the target organelle and distorted any pre-existing ER-membrane contact sites. The similar results at lipid droplets likely reflects that ER-lipid droplet contacts are more extensive than the discrete contacts at other organelles (Li et al., 2024). Given that the relocalization of LBR-FKBP-GFP can be used generically to mark different types of ER-MCS, we suggest that LaBeRling can be used as a multi-purpose label for ER-MCSs.

**Figure 5.**
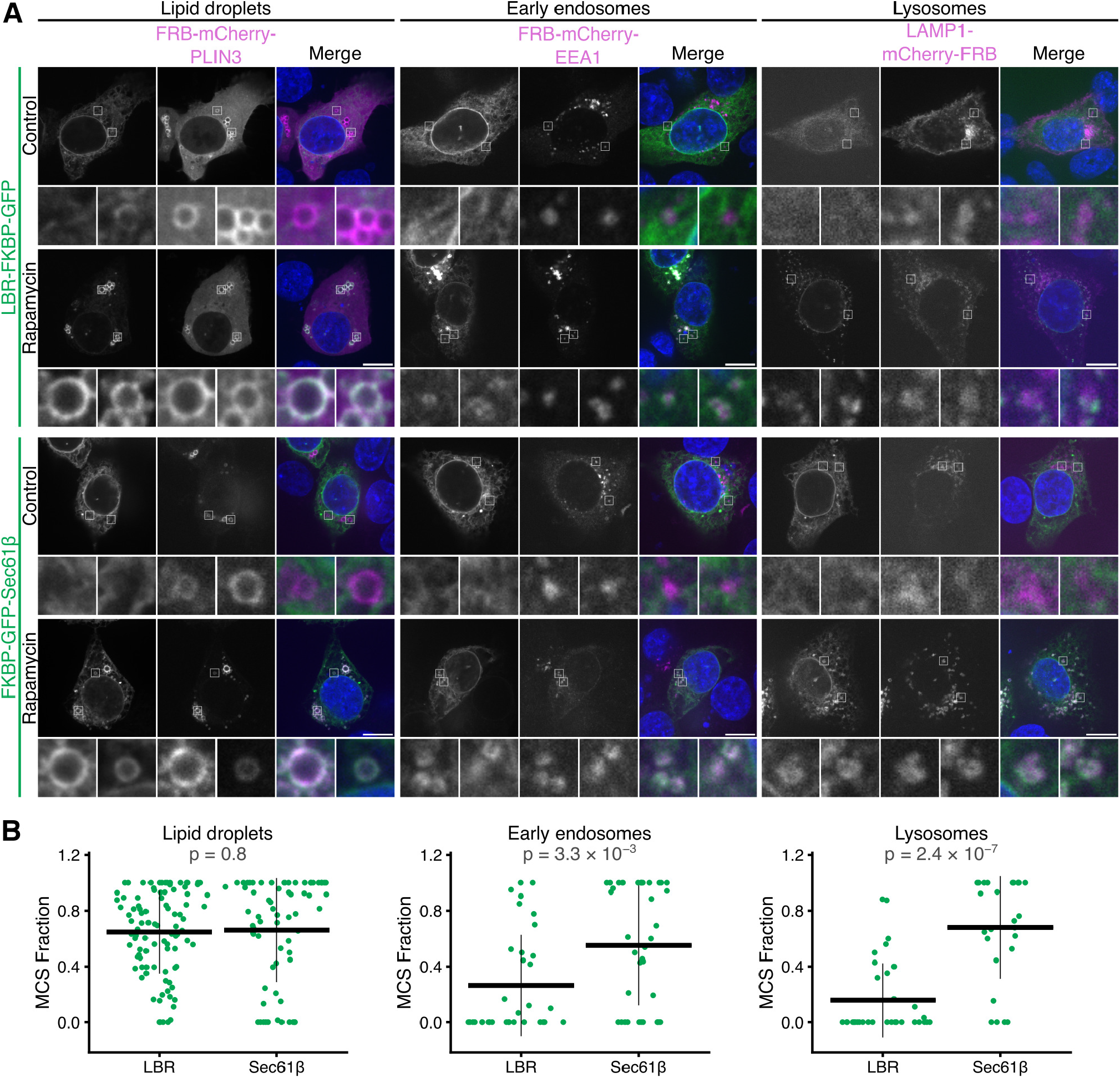
Using LBR-FKBP-GFP relocalization to mark ER-lipid droplet, ER-endosome or ER-lysosome contact sites. (**A**) Example micrographs of HCT116 cells expressing either LBR-FKBP-GFP or FKBP-GFP-Sec61β (green) together with the indicated mCherry-FRB tagged protein anchor localizing at the target membrane (magenta), either lipid droplets (FRB-mCherry-PLIN3), early endosomes (FRB-mCherry-EEA1), or lysosomes (LAMP1-mCherry-FRB), and stained with DAPI (blue). Lipid droplet number was increased by incubation with oleic acid (200 μM) for 17 h. Similar treatment reported to make no significant change to ER-Lipid droplet contacts (Valm et al., 2017). Relocalized samples were treated with rapamycin (200 nM) for 30 min before fixation. Control samples were not treated with rapamycin. A single slice from a z-stack of an interphase or metaphase cell are shown. Scale bars, 10 μm; zooms are 7.3× expansion of the ROI. (**B**) Plot of the fraction of the organelle perimeter that is above threshold. Quantification is of line profiles like those shown in Supplementary Figure S9. P-values, Tukey’s HSD *post hoc* test; n_cell_ = 4-29.

### LBR sterol reductase activity is not required for ER-MCS labeling

We wondered if LBR was unique in being able to be used in this way to label ER-MCSs. To look at this we investigated the relocalization of three other proteins: emerin, LAP2β and BAF; which each localize at least partially to the inner nuclear envelope. Each protein was tagged at the N-terminus with FKBP-GFP and expressed in HCT116 cells alone (control), or coexpressed with either Stargazin-mCherry-FRB or Mito-Trap, and all cells treated with rapamycin (200 nM). We found that each of the relocalized proteins completely coated the surface of the plasma membrane or mitochondria in interphase or mitosis (Supplementary Figure S10). This suggests that something unique to LBR meant that it can be used as an inducible ER-MCS marker.

LBR transmembrane regions are important for sterol reductase activity, essential in the cholesterol biosynthesis pathway (Tsai et al., 2016). To determine if the sterol reductase function is required to label MCSs, we generated two cholesterol synthesis point mutants (N547D or R583Q) associated with disease and tested their ability to inducibly label ER-PM MCSs in mitosis (Clayton et al., 2010). Both mutants were relocalized to discrete puncta that were indistinguishable from those formed after relocalization of the WT LBR-FKBP-GFP to Stargazin-mCherry-FRB (Figure 6A,B). We made a series of truncation constructs to try to isolate the minimal region needed for inducible labeling of ER-MCSs. The two smallest constructs of this series contained either the first two transmembrane (TM) domains or the first TM domain only (GFP-FKBP-LBR(1-288) or FKBPLBR(1-245)-GFP, respectively). Surprisingly, both of these constructs formed discrete clusters upon relocalization, similar to the full-length protein (GFP-FKBPLBR) (Figure 6B,C). These results suggest that the property of MCS targeting is contained in the N-terminal region and first TM domain; however further truncations of this region resulted in mislocalization of the construct from the ER. Like the point mutants, the truncated constructs are not predicted to have any sterol reductase activity, so we can rule out cholesterol biosynthesis as the reason why LBR can be used to label ER-MCSs.

**Figure 6.**
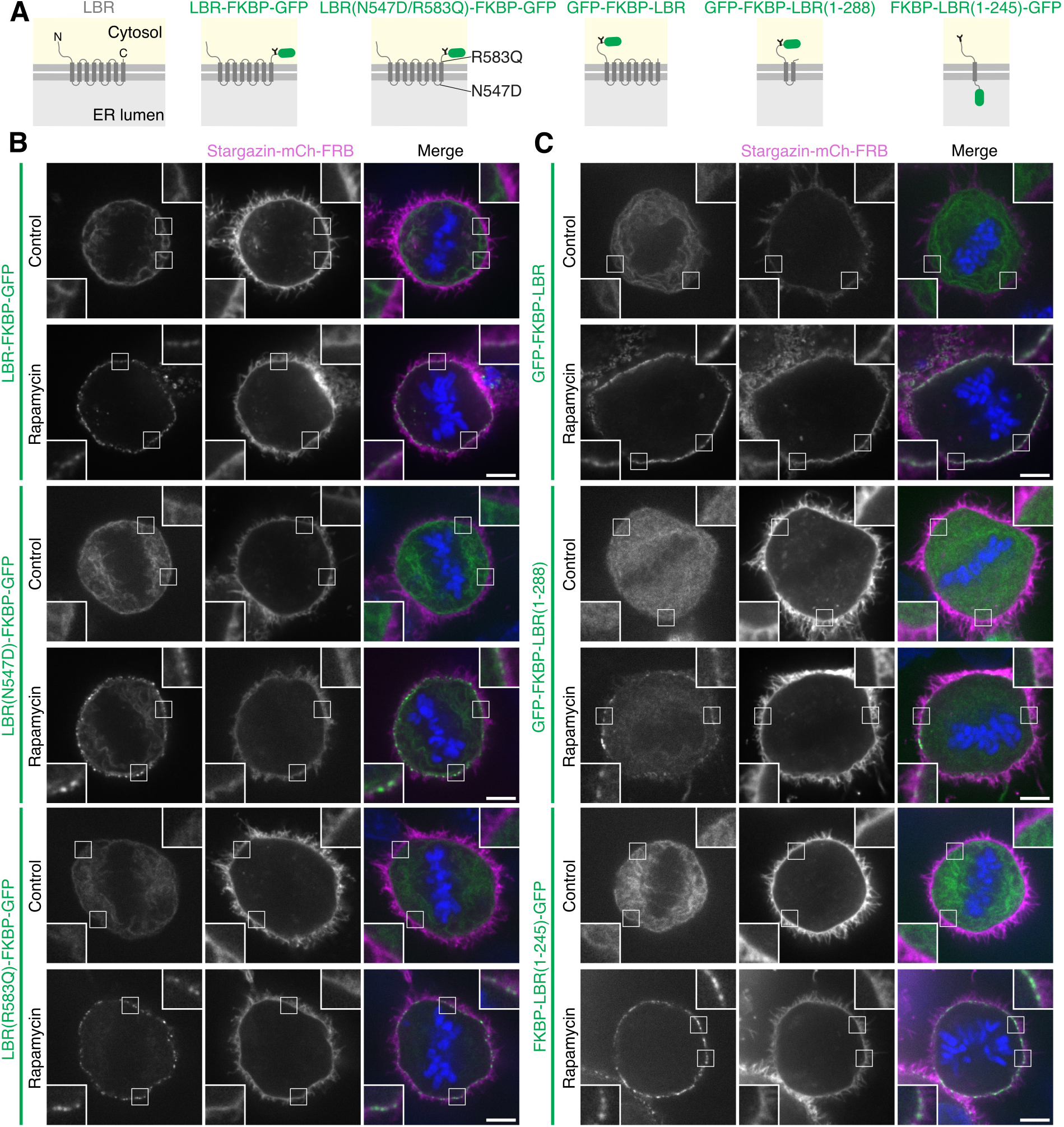
LBR cholesterol synthesis function is not required for labeling/truncated LBR has similar labeling to the full-length protein/LBR cholesterol synthesis mutants and truncated LBR form relocalized clusters. (**A**) Schematic of GFP- and FKBP-tagged LBR, sterol reductase point mutants (N547D or R583Q) and truncated proteins (expressing LBR amino acids 1-288 or 1-245). Example micrographs of HCT116 cells expressing FKBP- and GFP-tagged LBR, LBR sterol reductase mutants (**B**) or truncated LBR (**C**) (green) alongside Stargazin-mCherry-FRB (magenta), stained with DAPI (blue). Relocalized samples were treated with rapamycin (200 nM) for 30 min before fixation. Control samples were not treated with rapamycin. Scale bars, 5 μm; insets, 2.5× expansion of ROI.

Rather than a molecular determinant in the N-terminus of LBR, an alternative possibility is that expression level of LBR is such that the density of protein in the ER is sufficient to label contact sites, yet is not high enough to distort them. Western blot analysis indicated that the levels of expressed LBR-FKBP-GFP were indeed lower than FKBP-GFP-Sec61β (Supplementary Figure S11A,B). However, the expression of the truncated constructs GFP-FKBP-LBR(1-288) and FKBP-LBR(1-245)-GFP was higher than FKBP-GFP-Sec61β, and since these constructs could be used for LaBeRling, this argues against the idea that contact site labeling requires lower levels in the ER (Supplementary Figure S11A). We had already established that over-expression of LBR-FKBP-GFP results in LaBeRling of contact sites similarly to endogenously tagged LBR, so distortion of contact sites such as that seen with Sec61β and other proteins, is not possible to mimic with higher LBR expression. Therefore we sought to reduce the expression of FKBP-GFP-Sec61β to see if labeling of contact sites was possible. Even with lower expression, relocalization of FKBP-GFP-Sec61β to Stargazin-mCherry-FRB resulted in large contacts that were much larger than those marked with mScarlet-I3-6DG5-MAPPER (Supplementary Figure S11C and S12). These experiments suggest that simple expression differences are not sufficient to explain the difference in activity, and argue that LBR is especially suited for contact site labeling.

### Using LaBeRling to investigate novel contact sites: ER-Golgi contact sites in mitosis

Contact sites between the ER and *trans*-Golgi network (TGN) have been described (Venditti et al., 2019). The Golgi undergoes massive remodeling during mitosis (Carlton et al., 2020) but due to the lack of inducible labeling techniques, the status of ER-Golgi MCSs during mitosis has not been studied. Having established LaBeRling, we sought to establish whether we could label ER-Golgi MCSs and if so, to test if they persist during mitosis. HTC116 cells transiently co-expressing LBR-FKBP-GFP and FRB-mCherry-Giantin(3131-3259) were imaged following relocalization with rapamycin (200 nM) (Figure 7A). We compared these to similarly prepared samples co-expressing FKBP-GFP-Sec61β and FRB-mCherry-Giantin(3131-3259). In interphase cells, LBR-FKBP-GFP formed distinct puncta when relocalized to FRB-mCherry-Giantin(3131-3259) Golgi structures. By contrast, FKBP-GFP-Sec61β coated much of the interphase Golgi structures after relocalization (Figure 7A,B). Similar LaBeRling was also observed in RPE-1, Cos-7 and HeLa cells (Supplementary Figure S6). These results suggest that LBR can be used to selectively label ER-Golgi MCSs.

**Figure 7.**
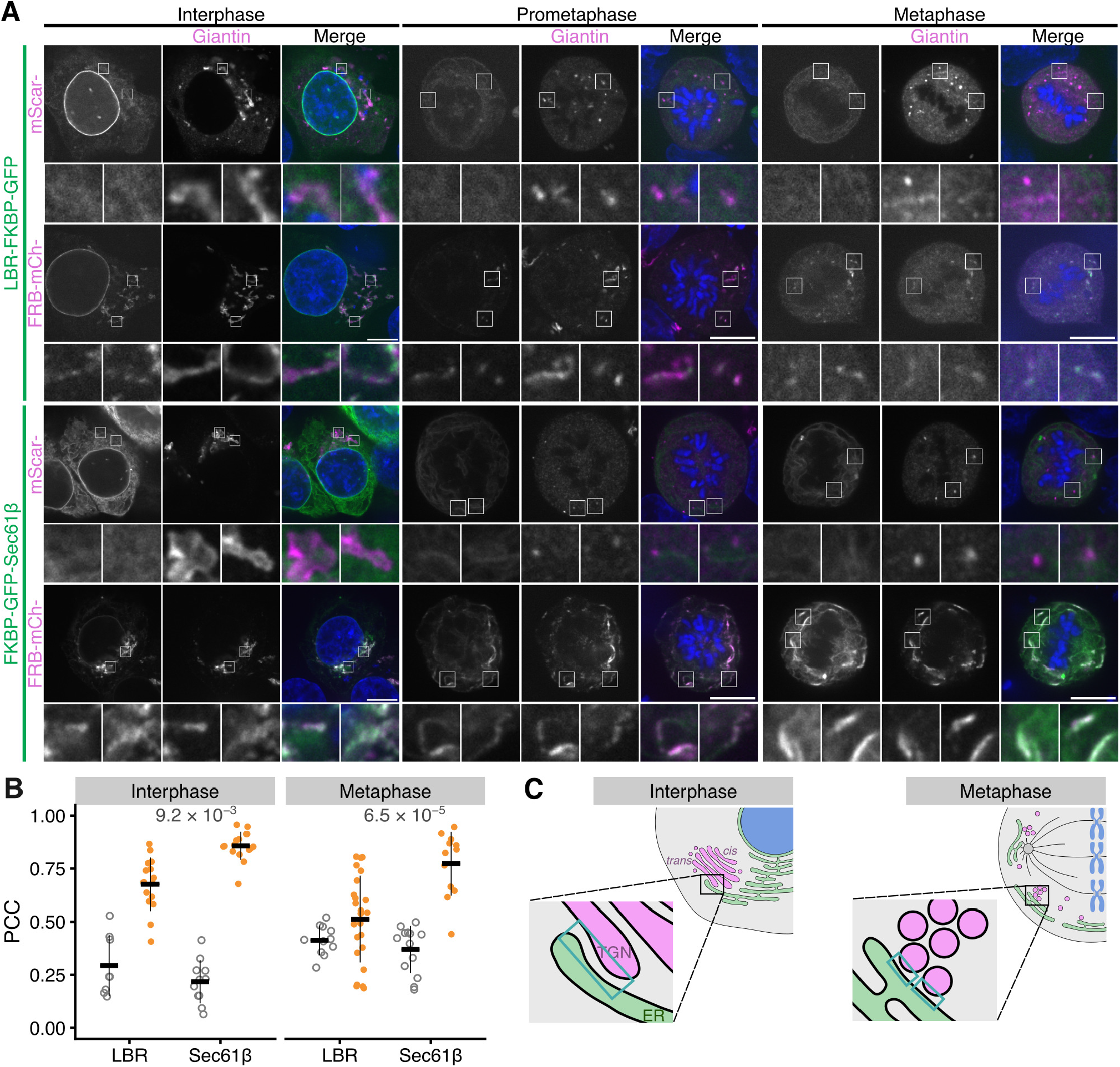
Using LBR-FKBP-GFP relocalization to identify novel ER contact sites. (**A**) Example micrographs of synchronized HCT116 cells expressing either LBR-FKBP-GFP or FKBP-GFP-Sec61β (green) together with mScarlet-Giantin or FRB-mCherry-Giantin(3131-3259) (magenta), and stained with DAPI (blue). All samples were treated with rapamycin (200 nM) for 30 min before fixation. Single slices from z-stacks of an interphase, early mitotic or metaphase cell are shown. Scale bars, 10 μm; zooms, 5.8× expansion of ROI for interphase cells and 4× expansion for prometaphase and metaphase. (**B**) Plots of Pearson’s corelation coefficient (PCC) between LBR-FKBP-GFP or FKBP-GFP-Sec61β and either mScarlet-Giantin (gray) or FRB-mCherry-Giantin(3131-3259) (orange) at the indicated cell cycle stage. Markers, cells. Bars indicate mean ± sd. P-values, Tukey’s HSD *post hoc* test; n_cell_ = 8-24. (**C**) Schematic representation of ER-Golgi contacts in interphase or metaphase cells, with example contact sites indicated in the expanded region (blue box).

In mitotic cells, relocalized LBR-FKBP-GFP was observed to be at puncta coinciding with Golgi fragments at prometaphase and at metaphase (Figure 7A). This relocalization pattern could be observed in live mitotic cells (SV8) forming with dynamics similar to that of other ER-MCS labeling events. Moreover, in mitotic HCT116 LBR-FKBP-GFP knock-in cells the same labeling was present confirming that the labeling of ER-Golgi MCSs in mitosis was not due to overexpression of the LBR protein (Figure 8A). Again, by contrast, large patches of FKBP-GFP-Sec61β signal were observed at relocalized at FRB-mCherry-Giantin(3131-3259) Golgi fragments in prometaphase and metaphase cells (Figure 7A,B). In fact, the Golgi haze appeared to be cleared from the cytoplasm and that Golgi was recruited to the ER. These results suggest that selective labeling of ER-Golgi MCSs by LBR was possible and was distinguishable from non-specific heterodimerization between ER and Golgi.

**Figure 8.**
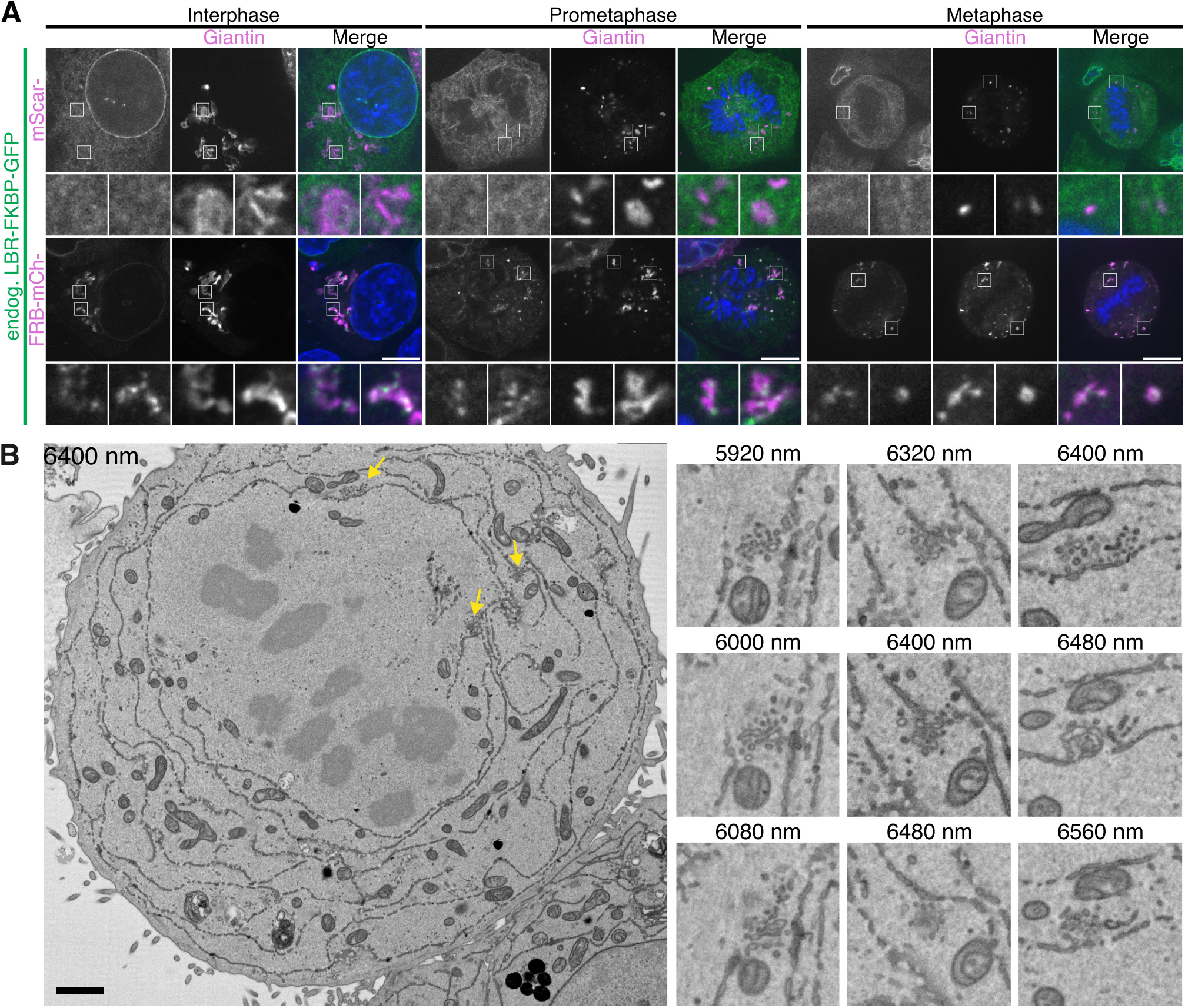
LaBeRling ER-Golgi membrane contact sites in mitosis. (**A**) Example micrographs of synchronized HCT116 LBR-FKBP-GFP knock-in cells expressing either mScarlet-Giantin or FRB-mCherry-Giantin(3131-3259) (magenta), and stained with DAPI (blue). All samples were treated with rapamycin (200 nM) for 30 min before fixation. Single slices from z-stacks of an interphase, early mitotic or metaphase cell are shown. Scale bars, 10 μm; insets, 4× expansion of ROI. (**B**) Single slices from SBF-SEM dataset of a metaphase HCT116 LBR-FKBP-GFP knock-in cell transiently expressing Mitotrap and treated with rapamycin (200 nM) are shown. Depth of each slice within the dataset (nm) is indicated. Example Golgi clusters are shown by yellow arrows on the full slice image. Three sequential slices of these regions (3× expansion) are shown beside. Scale bars, 2 μm and 0.5 μm on zoom region.

The persistence of LaBeRling in mitotic cells suggests that ER-Golgi MCSs are maintained during mitosis. To examine these contacts in further detail, we examined ER-MCSs in mitotic HCT116 LBR-FKBP-GFP knock-in cells by SBF-SEM (Figure 8B and Supplementary Figure S13). Small clusters of vesicles were readily observable within 30 nm of ER in metaphase and telophase cells. These clusters match the mitotic Golgi clusters described in EM images of HeLa cells in prometaphase, metaphase and telophase (Lucocq et al., 1987), and more recently in NRK cells in prophase, metaphase and late anaphase (Jokitalo et al., 2001). Together these data suggest that ER-Golgi MCSs are maintained in mitotic cells and can be labeled by the relocalization of LBR-FKBP-GFP to Golgi membranes using FRB-mCherry-Giantin(3131-3259).

## Discussion

In this study, we developed a new method, LaBeRling, for inducible labeling of ER-MCSs using the ER protein LBR. Unlike previous inducible labeling approaches, LaBeRling does not alter existing ER-plasma membrane or ER-mitochondria MCSs and it does not induce artificial contacts. Moreover, labeling is fast (<2 min) and persists over many minutes without distortion of MCSs, so it is ideal for labeling MCSs at discrete stages of the cell cycle. Finally, we used this method to demonstrate the presence of ER-Golgi MCSs in mitosis, at a time of mitotic Golgi dispersal.

LaBeRling uses heterodimerization of LBR with a generic anchor protein on the target membrane. This anchor protein does not need to localize to MCSs, and can be readily interchanged, so that LBR can be used to label a range of ER-MCSs (between plasma membrane, mitochondria, early endosomes, lysosomes, lipid droplets, and Golgi) in interphase and mitotic cells. In the past, other approaches have used MCS tethering proteins for inducible labeling (Chung et al., 2015; Lees et al., 2017), with the logic that this will increase specificity. This is problematic because both the overexpression of tethering proteins themselves and their subsequent heterodimerization can distort MCSs and induce artificial contacts (Chung et al., 2015; He et al., 2017), as similarly seen using membrane targeting anchors (Csordás et al., 2010; Komatsu et al., 2010; Miner et al., 2024). LaBeRling delivers more universal labeling of contact sites since the initial coverage of the target membrane and the ER is diffuse and homogeneous yet specific to the respective compartments. We envisage that LaBeRling could also be applied to label other ER-membrane contacts, for example, ER-peroxisome and ER-autophagosome MCSs.

Our serendipitous discovery that LBR can be used to label MCSs is intriguing: what makes LBR so special? We saw that the anchor protein remains homogeneously distributed after LBR relocalization and so the MCS specificity of the labeling seems to be driven by LBR, from the ER side. This observation could mean that ER-MCSs are not truly bipartite and is supported by work showing that altered diffusivity of an ER-side contact site protein VAPB can affect ER-mitochondria MCSs (Obara et al., 2024). Why LBR behaves in this way, whereas other ER proteins do not, is unresolved. Cholesterol synthesis activity was not essential for LaBeRling because point mutants and truncated LBR proteins that lack sterol reductase activity all labeled EM-plasma membrane MCSs similarly to fulllength WT LBR. It is possible that the intermembrane distance at the ER-MCSs is such that the size of LBRFKBP-GFP is in a “sweet spot” to be immobilized by the heterodimerization procedure, but not so large as to crosslink membranes outside of the MCS. In support of this possibility there is evidence that altering the spacing in optogenetic heterodimerization of ER-plasma membrane linkages affects labeling efficiency (He et al., 2017). Adjusting linkage distance of ER-plasma membrane contacts has also been indicated to affect protein translocation into the MCS (Várnai et al., 2007; Chang et al., 2013). However, the intermembrane distances of MCSs are reported to be rather variable and we observed that various anchor proteins can be used successfully, which argues against the idea that physical spacing can explain the labeling behavior. A further possibility is that the density of LBR in the ER is lower than other proteins so that its relocalization doesn’t distort contacts. However, we saw no distortion at higher expression levels, and also that reducing the expression of other ER proteins didn’t allow for specific labeling, which argues against this possibility. Recent work points to LBR having affinity for the IP_3_ receptor, which could potentially provide a molecular explanation for why it can be used to inducibly label contacts (Zhao et al., 2024). Whatever the mechanism, the observation that sterol reductase activity is not required means that the LBR point mutants can be used to ensure the local sterol environment at the MCS is not modified by labeling. This may be important because MCSs are reported to be enriched with cholesterol (Hayashi and Fujimoto, 2010; Fujimoto et al., 2012; Zung et al., 2024), and an ideal labeling method would not perturb the lipids at the endogenous MCS. Conversely, additional protein domains could be added to LBR to deliver new enzymatic activities to ER-MCSs in order to experimentally manipulate these areas and answer new biological questions.

In interphase, ER-TGN MCSs have been described (Wu et al., 2018; Venditti et al., 2019) and we used a generic Golgi anchor protein and LBR-FKBP-GFP to detect ERGolgi MCSs, which were presumably ER-TGN contacts. We used our inducible method to show that ER-Golgi MCSs are maintained in mitosis, a time when the Golgi has been disassembled (Carlton et al., 2020). To our knowledge this is the first labeling of these MCSs during mitosis but is corroborated by evidence from electron microscopy studies. Briefly, the Golgi is disassembled by severing the Golgi ribbons into stacks and then dispersing the stacks into Golgi “blobs” and “haze” (Misteli and Warren, 1995), which correspond to vesicular clusters and single vesicles, respectively. Each Golgi cluster likely contains a mixture of vesicles from TGN, *cis*- and medial-Golgi. For example, a subset of the vesicles within the mitotic Golgi clusters had TGN markers in HeLa cells by EM (Lucocq et al., 1987) and an incomplete overlap of different Golgi markers stained within mitotic Golgi clusters was detected by light microscopy in NRK cells (Puri et al., 2004). Therefore ER-TGN contacts are an efficient way for the ER to remain in contact with Golgi clusters. The function of ER-Golgi contacts maintained during mitosis is unclear. The spindle operates in an “exclusion zone” which is largely membrane free (Nixon et al., 2017). Maintained ER-Golgi contacts could serve simply to exclude clusters from the spindle area and to prevent these membrane fragments from interfering with chromosome segregation. Another possibility is that the ER-Golgi contacts may coordinate reassembly by allowing the clusters to surf on the ER towards the spindle pole and the midbody where the Golgi twins coalesce and begin to reassemble (Gaietta et al., 2006).

We used LaBeRling to probe other ER-MCSs in mitosis. For example, we saw that the total number of ER-plasma membrane MCSs was reduced at metaphase compared with interphase cells. This observation is seemingly at odds with our observation that the density of contacts remained constant. However, our density measurement uses a surface constructed from the contact sites rather than the plasma membrane surface area for normalization. So while the net number of contacts decreases, the spacing between them is maintained as the plasma membrane is drawn up and away from the underlying ER (Erickson and Trinkaus, 1976). How this process is regulated and how MCS are modified during mitosis are interesting questions for the future.

To conclude, the method we describe to label ER-MCS using LBR can be applied to a wide variety of questions at many different types of contact in cells. It can be further developed to tweak the properties of ER-MCSs or be engineered to allow reversible highlighting of membrans contact sites.

## Methods

### Molecular biology

The following plasmids were available from Addgene or previous work: pEBFP2-N1 (Addgene #54595); EGFP-BAF (Addgene #101772); Emerin pEGFP-C1 (637) (Addgene #61993); FKBP-alpha(740-977)-GFP (Addgene #100731); FKBP-GFP-Sec61β (Addgene #172442); FRB-mCherry-Giantin (Addgene #186575); GFP-EEA1 (Addgene #42307); LAMP1-mCherry-FRB (Addgene #186576); LAP2 Full I pAcGFP-N1 monomeric GFP (1317) (Addgene #62044); LBR pEGFP-N2 (646) (Addgene #61996); pFKBP-GFP-C1 (Clarke and Royle, 2018); pMaCTag-P05 (Addgene #120016); pMito-mCherry-FRB (Addgene #59352); pMito-mCherry-FRB(T2098L); pMito-dCherry-FRB (Addgene #186573); pmScarlet-Giantin-C1 (Addgene #85048); SH4-FRB-mRFP (Addgene #100741); Stargazin-dCherry-FRB (Addgene #172444); Stargazin-GFP-LOVpep (Addgene #80406); Stargazin-mCherry-FRB (Addgene #172443) (Ferrandiz et al., 2022; Wood et al., 2017; Küey et al., 2022; Fesenko et al., 2024).

To generate the pFRB-mCherry-C1 vector, FRB was amplified from pMito-mCherry-FRB plasmid (using CG CGGCTAGCGGCCACCATGATCCTCTGGCATGAGATGTGGCATGAAGGC and TCGCaccggtggGCCGGCC TGCTTTGAGATTCGTCGGAACAC) and inserted into pmCherry-C1 vector (using NheI-AgeI). pmCherry-C1 vector was made by substituting mCherry for EGFP in pEGFP-C1 [Clontech] by AgeI-XhoI digestion.

LBR-FKBP-GFP was generated by cutting LBR from LBR-mCherry using BamHI-KpnI sites and ligating into pFKBP-GFP-N1 plasmid, where FKBP was inserted into pEGFP-N1 at BamHI-AgeI sites. Similarly, FKBP-GFP-emerin was made by digestion of Emerin pEGFP-C1 (637) and ligation into FKBP-GFP-C1 plasmid using XhoI and BamHI sites. To clone FKBP-GFP-LAP2β, BglII-SalI sites were introduced at either end and a C-terminal stop codon to LAP2β amplified by PCR from LAP2 Full I pAcGFP-N1 monomeric GFP (1317) plasmid template (primers aagcttAGATCTATGCCGGAGT TCCTAGAGG and tcgagGTCGACCTAgCAGTTGGA TATTTTAGTATCTTGAAG), and ligating into pFKBP-GFP-C1. FKBP-GFP-BAF was generated by digestion of mCherry-BAF plasmid and ligation into pFKBP-GFP-C1 at BglII and Acc65I sites. The mCherry-BAF plasmid used in this cloning was made by PCR amplification of the BAF-encoding region from EGFP-BAF (primers aag cttAGATCTATGACAACCTCCCAAAAGC and tcgagAA GCTTCTACAAGAAGGCATCACACC) and ligation into pmCherry-C1 vector (BglII-HindIII)

LBR point mutations (N547D or R583Q) were introduced by site-directed mutagenesis using LBR-FKBP-GFP plasmid template. Primer with mismatches were designed to introduce the mutation N547D (GTTCG CCACCCCGATTACTTGGGTGATCTCATC and GA TGAGATCACCCAAGTAATCGGGGTGGCGAAC) or R583Q (CATGTTGCTTGTCCACCAAGAAGCTCGTG ACG and CGTCACGAGCTTCTTGGTGGACAAGCAA CATG).

The LBR truncations were generated by PCR amplication of the LBR region from LBR pEGFP-N2 (646) template. To generate the GFP-FKBP-LBR(1-288) plasmid, the primer set aagacaGAATTCaATGCCAAGTA GGAAATTTGCCG and tcgagGGATCCTTAATCAATA AGAGGCGTTCCTTCTACAAC was used. The PCR product was ligated into pEGFP-FKBP-C1 using EcoRI-BamHI sites. FKBP-LBR(1-245)-GFP truncation was made by amplifying the region encoding amino acids 1-245 (using aagacaGGTACCGTGAACCGTCAGATCCG CTAG and tcgagACCGGTagAGGAGGGAAATTCAGAAGACTGGGATCTTTC). The PCR product was ligated in substitution of the AP2A1 encoding region in FKBP-alpha(740-977)-GFP (using KpnI-AgeI).

The plasma membrane anchor Stargazin-EBFP2-FRB was made by PCR of Stargazin encoding region from Stargazin-GFP-LOVpep (primers gcggctagcATGGGG CTGTTTGATCGAGGTGTTCA and TTTACTCATGG ATCCttTACGGGCGTGGTCCGG) and then ligating into pEBFP2-N1. Stargazin-EBFP2 was amplified from the resulting plasmid (primers aagcttGCTAGCcATGG GGCTGTTTGATCGAGG and tcgagGGTACCccCTTG TACAGCTCGTCCATGC), excluding the stop codon, and ligated N-terminal to FRB in a plasmid encoding Stargazin-dCherry-FRB. SH4-FRB-EBFP2 was made by substituting mRFP of SH4-FRB-mRFP with EBFP2 from pEBFP2-N1 (using AgeI-NotI).

Endosomal anchor FRB-mCherry-EEA1 was made by cutting full-length EEA1 from GFP-EEA1 using XhoI-BamHI and ligating into pFRB-mCherry-C1. Lipid droplet anchor FRB-mCherry-PLIN3 was made from a mGFP-PLIN3 full-length plasmid that was generated by DNA synthesis and Gibson Assembly, cutting FRB-mCherry from FRB-mCherry plasmid (NheI-BsrGI) and substituting for the mGFP of mGFP-PLIN3.

Plasma membrane anchor binding to rapalog, Stargazin-mCherry-FRB(T2098L), was made by cutting the Stargazin-encoding sequence from Stargazin-mCherry-FRB plasmid (NheI-BamHI) and paste in place of the mitochondrial targeting sequence in pMito-mCherry-FRB(T2098L).

The template plasmid for the C-terminal PCR tagging CRISPR method (Fueller et al., 2020) encoding FKBP-GFP tag (pMaCTag-P05-FKBP-GFP) was generated by amplifying the region encoding FKBP-GFP from pFKBP-GFP-N1 and introducing BamHI-SpeI restriction sites (primers aagcttGGATCCCCGCCACCAA TGGGAGTGCAGGTGG and tcgagACTAGTTTACTTG TACAGCTCGTCCATGCCGAGAGT), and then ligating in place of the GFP tag in available pMaCTag-P05 to give pMaCTag-P05 FKBP-GFP. To generate the PCR product for editing, LBR-specific tagging oligos were designed using the online design tool (M1, CGTGACGAG TACCACTGTAAGAAGAAATACGGCGTGGCTTGGGAAAAGTACTGTCAGCGTGTGCCCTACCGTATATTTCCATACATCTACTCAGGTGGAGGAGGTAGTG and M2, TTTGCAAATGGCAGCTGGAATTGCAGGAGTATTTTGTAGAAAAGCCAGAAGAGCAAAAAAAAGAGCATTAGTAGATGTATGATCTACACTTAGTAGAAATTAGCTAGCTGCATCGGTACC). Primers for genotyping PCR to confirm LBR-FKBP-GFP knock-in were AGAAT TTGGGGGAAAGCAGG and CATCCTTACTTGTATTTTTCCTATGTTAACTG or AAGACAATAGCAGGCATG CT and CAGTGGCACCATAGGCATAA.

The mScarlet-I3-6DG5-MAPPER plasmid was designed based on the available GFP-MAPPER sequence and generated by DNA synthesis (Chang et al., 2013). Briefly, the GFP-encoding sequence of GFP-MAPPER was replaced with mScarlet-I3 and the unwanted FRB sequence was substituted with a codon-optimized sequence of Neoleukin-2/15 (PDB code, 6DG5), which is structurally similar and close in mass to FRB. Linker sequence lengths were maintained similar to that in the GFP-MAPPER sequence.

### Cell biology

HCT116 (CCL-247; ATCC) cells and lines derived from HCT116 were maintained in DMEM supplemented with 10 % FBS and 100 U mL^−1^ penicillin/streptomycin. All cell lines were kept in a humidified incubator at 37 °C and 5 % CO_2_. Cells were routinely tested for mycoplasma contamination by a PCR-based method.

HCT116 LBR-FKBP-GFP CRISPR knock-in cells were generated using a C-terminal PCR tagging CRISPR method (Fueller et al., 2020). HCT116 cells were transfected with the M1-M2 PCR cassette and a plasmid encoding Cas12a (pVE13300). Cells were selected and maintained in media supplemented with puromycin dihydrochloride (Gibco) at 1.84 μM. Populations of cells positive for GFP signal were selected by FACS and the positive pools of cells were characterized by genotyping PCR, western blot and microscopy.

For transient transfection, Fugene-HD (Promega) was used to transfect HCT116 and RPE-1 cells, and Gene-Juice (Merck) used for HeLa and Cos-7 cells, each according to the manufacturer’s instructions. A total of 1 μg DNA with 3 μL reagent was used per fluorodish or well of a six-well plate for each transfection reagent. With the exception of LBR-FKBP-GFP expressed endogenously, the expression of all constructs was via transient transfection.

Heterodimerization of FKBP and FRB tags was induced through addition of rapamycin to media at a final concentration of 200 nM. For fixed cell experiments, 200 nM rapamycin (J62473, Alfa Aesar) was prepared in complete media 2 mL per well of a 6 well plate; growth media was removed, rapamycin-containing media added, and then the plate was returned to the incubator until fixation. To apply rapamycin to live cells on the microscope, rapamycin solution (400 nM) in imaging media (Leibovitz’s L-15 Medium, no phenol red, supplemented with 10 % FBS) was diluted (1:2) to final concentration 200 nM using media in the dish. Similarly, heterodimerization of FKBP and FRB(T2098L) tags was induced with rapalog (A/C Heterodimerizer, Takara, 635057) at a final concentration of 5 μM, applied to cells similarly to the rapamycin treatment described above. To visualize DNA in live cells, dishes were incubated for 15–30 min with 0.1 μM SiR-DNA (Spirochrome) prepared in complete media. Cells were selected for moderate expression of the anchor construct, to minimize the possibility that organelle morphology, and MCSs, were affected by each anchor.

Cells were synchronized by treatment with thymidine (2 mM, 16 h), followed by washout, 7–8 h incubation and then RO-3306 treatment (9 μM, 16 h). Cells were released from synchronization and incubated for 15 min or 30 min before applying rapamycin solution (final concentration 200 nM) and incubating a further 30 min before fixation.

To increase the number of lipid droplets, the media on cells was supplemented with oleic acid (O3008, Sigma) at 200 μM) for around 17 h before fixation.

Thapsigargin treatment was used to induce an increase in ER-plasma membrane contacts, as described (Orci et al., 2009; Chang et al., 2013). Thapsigargin (T9033, Sigma) solution was prepared at 1.5 mM in DMSO and applied to cells at 1 μM final concentration in imaging media.

### Fluorescence methods

For fixed-cell imaging, cells were seeded onto glass cover slips (16 mm diameter and thickness number 1.5, 0.16–0.19 mm). Cells were fixed using 3 % PFA/4 % sucrose in PBS for 15 min. After fixation, cells were washed with PBS and then incubated in permeabilization buffer (0.5 % v/v Triton X-100 in 1xPBS) for 10 min. Cells were washed twice with PBS, before 45–60 min blocking (3 % BSA, 5 % goat serum in 1xPBS). Antibody dilutions were prepared in blocking solution. After blocking, cells were incubated for 1 h with primary antibody, PBS washed (three washes, 5 min each), 1 h secondary antibody incubation, PBS washed (three washes, 5 min each), mounted with Vectashield containing DAPI (Vector Labs Inc.) and then sealed. PhenoVue Fluor 568 – Concanavalin A dye (CP95681) stain was added to coverslips at 50 μg mL^−1^ in HBSS for 15 min at room temperature, followed by three 5 min PBS washes before mounting and sealing as described above.

### Western blotting

For Western blotting, cells were harvested, and lysates were prepared by sonication of cells in UTB buffer (8 M urea, 50 mM Tris, and 150 mM 2-mercaptoethanol) for HCT116 LBR-FKBP-GFP knock-in cell characterization (Figure S2). For collection of lysates for comparison of expression levels (Figure S11), lysis buffer containing 50 mM Tris-HCl, pH 7.0 (at 25 °C), 150 mM NaCl, 1 mM EDTA, 1 mM dithiothreitol (DTT) was used. All lysates were incubated on ice for 30 min, clarified in a benchtop centrifuge (20 800 *g*) for 15 min at 4 °C, boiled in Laemmli buffer for 10 min, and resolved on a precast 4–15 % polyacrylamide gel (Bio-Rad). Proteins were transferred to nitrocellulose using a TransBlot Turbo Transfer System (Bio-Rad) (blots shown in Figure S2) or iBlot 2 Dry Blotting System (Invitrogen, IB21001) (blots in Figure S11). The following antibodies were used: rat monoclonal anti-GFP (3h9, Chromotek) at 1:1000 in 3 % BSA TBST (Figure S2); rabbit anti-GFP (ab6556, Abcam) at 1:1000 in 2 % milk TBST (Figure S11); mouse anti-LBR antibody (polyclonal) (SAB1400151, Sigma-Aldrich) at 1:500 in 2 % milk TBST; goat anti-rat IgG-Peroxidase antibody (A9037, Sigma) at 1:5000 in 5 % milk TBST; sheep anti-mouse IgG-HRP (NXA931, Cytiva) at 1:5000 in 5 % milk TBST; donkey anti-rabbit IgG-HRP (NA934V, Cytiva) at 1:2500 in 2 % milk TBST; loading control HRPconjugated mouse anti-β-actin (C4) (sc-47778, Santa Cruz Biotechnology) at 1:20000 in 2 % milk TBST (Figure S2) or mouse anti-β-actin (A2228, Sigma) with sheep anti-mouse IgG-HRP (NXA931, Cytiva) used at 1:2000 in 2 % milk TBST (Figure S11).

### Microscopy

As described previously, images were captured using a Nikon CSU-W1 spinning disc confocal system with SoRa upgrade (Yokogawa) with either a Nikon, 100×, 1.49 NA, oil, CFI SR HP Apo TIRF or a 63×, 1.40 NA, oil, CFI Plan Apo objective (Nikon) with optional 2.8× intermediate magnification and 95B Prime camera (Photometrics) (Ferrandiz et al., 2022). The system has a CSU-W1 (Yokogawa) spinning disk unit with 50 μm and SoRa disks (SoRa disk used), Nikon Perfect Focus autofocus, Okolab microscope incubator, Nikon motorized xy stage and Nikon 200 μm z-piezo. Excitation was via 405 nm, 488 nm, 561 nm and 638 nm lasers with 405/488/561/640 nm dichroic and Blue, 446/60; Green, 525/50; Red, 600/52; FRed, 708/75 emission filters. Acquisition and image capture was via NiS Elements (Nikon). All microscopy data was stored in an OMERO database in native file formats.

### Serial block face-scanning electron microscopy

Preparation of samples for serial block face-scanning electron microscopy (SBF-SEM) was performed as described previously (Ferrandiz and Royle, 2023; Ferrandiz et al., 2022). Briefly, HCT116 LBR-FKBP-GFP knock-in cells expressing MitoTrap were plated onto gridded dishes and prior to imaging, incubated for around 30 min with 0.5 μM SiR-DNA (Spirochrome) to visualize DNA. Using light microscopy, live cells were imaged to confirm the induced labeling following rapamycin (200 nM) treatment. Control cells not treated with rapamycin were imaged in parallel and the coordinate position of the cell of interest recorded for correlation by SBF-SEM. Cells were washed twice with PB (phosphate buffer) before fixing (2.5 % glutaraldehyde, 2 % paraformaldehyde, 0.1 % tannic acid (low molecular weight) in 0.1 M phosphate buffer, pH 7.4) for 1 h at room temperature. Samples were washed three times with PB and then post-fixed in 2 % reduced osmium (equal volume of 4 % OsO_4_ prepared in water and 3 % potassium ferrocyanide in 0.1 M PB solution) for 1 h at room temperature, followed by a further three washes with PB. Cells were then incubated for 5 min at room temperature in 1 % (w/v) thiocarbohydrazide solution, followed by three PB washes. A second osmium staining step was then included, incubating cells in a 2 % OsO_4_ solution prepared in water for 30 min at room temperature, followed by three washes with PB. Cells were then incubated in 1 % uranyl acetate solution at 4 °C overnight. This was followed by a further three washes with PB. Walton’s lead aspartate was prepared adding 66 mg lead nitrate (TAAB) to 9 mL 0.03 M aspartic acid solution at pH 4.5, and then adjusting to final volume of 10 mL with 0.03 M aspartic acid solution and to pH 5.5 (pH adjustments with KOH). Cells were incubated in Walton’s lead aspartate for 30 min at room temperature and then washed three times in PB. Samples were dehydrated in an ethanol dilution series (30 %, 50 %, 70 %, 90 %, and 100 % ethanol, 5 min incubation in each solution) on ice, then incubated for a further 10 min in 100 % ethanol at room temperature. Finally, samples were embedded in an agar resin (AGAR 100 R1140, Agar Scientific). SBF-SEM imaging was carried out by the Biomedical Electron Microscopy Unit at University of Liverpool, UK.

## Data analysis

Analysis of confocal z-stacks of LBR-FKBP-GFP clusters at the plasma membrane was by 3D Spot Finder in 3D Image Suite plugin in Fiji. Briefly, outputs were fed into R where the size and number of clusters was stored. Contact area was defined as half of the surface area of a cluster. The location of clusters was used to find the surface of the cell using alphashape3d. The total number of clusters divided by the surface area of the alpha shape was used to determine the density of clusters per cell. For analysis of cluster formation in movies of LBR-FKBP-GFP relocalization to the plasma membrane, a weka segmentation method was used in TrackMate/Fiji to define the clusters that formed and track individual clusters over time. The outputs of these TrackMate XML files was analyzed using TrackMateR (Sittewelle and Royle, 2024). Tracks shorter than 4 frames or those that terminated before 100 s were removed from analysis and the remainder analyzed for shape, intensity, and number over time.

Analysis of ER-PM contacts labelled by LBR-FKBP-GFP/MAPPER was done using a semi-automatated procedure to segment each channel in 3D using 3D spot finder in 3D Image Suite in Fiji. Output were processed in R, where the total numbers of clusters per cell was analyzed per cell. Comparisons between pre and post rapamycin was done using paired t-tests on the individual cell data, with Holm-Bonferroni correction for multiple testing. Comparison between conditions was done using the experimental means using ANOVA with Tukey’s post hoc test.

Line profile data was harvested by manually outlining organelles of interest in the red channel, without sight of the green channel. Intensities for both channels along the profile were saved along with the background and maximum value for the image for normalization. Following import into R, the rle function was used to quantify segmented regions of the profile above a threshold.

For ER-mitochondria contact analysis, each SBF-SEM dataset was aligned using SIFT and cropped. A subvolume of one dataset (48 slices) was manually segmented for ER and mitochondria using IMOD. The segmentation and corresponding raw data were used to train nnU-Net v2 (Isensee et al., 2021) running on a GPU workstation (Intel Core i9-7900X, 128 GB, with TITAN Xp GPU). The resulting model was used to infer the ER and mitochondria in all datasets. Visualization of the inference maps or the manual segmented IMOD models was done using ChimeraX. A series of Fiji/ImageJ scripts were used to process the output. Briefly, an exact euclidean distance transform (EDT) was generated using the mitochondria channel to give a 3D volume of distances from each voxel to the nearest mitochondrion. Then the overlap between ER channel and EDT at distances of 10– 50 nm was calculated in 10 nm increments, to give the ER regions (contacts) within the appropriate distance from the mitochondrion. This result was segmented in 3D to classify all of the contacts. In addition, the mitochondria channel was also segmented in 3D. These outputs were processed in R to match each contact with its corresponding mitochondrion, which allowed the comparison of total contact surface area per mitochondrion with the mitochondrion surface area.

## Data and software availability

All code used in the manuscript is available at https://github.com/quantixed/p65p038 and additional data is available at Zenodo (https://doi.org/10.5281/zenodo.11396014).

## ACKNOWLEDGEMENTS

We thank Maelle Lorvellec and Laura Cooper from the Computing and Advanced Microscopy Unit (CAMDU) for their help and support. Alex Moore provided technical assistance to make the LBR-FKBP-GFP knock-in cells and to manually segment SBF-SEM data for nnU-Net training. Alison Beckett at the Liverpool Biomedical EM Unit carried out SBF-SEM imaging. We also acknowledge the help of Steven Servin-Gonzalez and the use of the Flow Cytometry Shared Resource Laboratory at the University of Warwick. We are grateful to members of the Royle lab for feedback on the project and manuscript. This work was supported by a Programme Award from Cancer Research UK (C25425/A27718). LD and MJ were supported by studentships from BBSRC Midlands Integrative Biosciences Training Partnership (MIBTP2, BB/M01116X/1 and MIBTP2020, BB/T00746X/1).

## AUTHOR CONTRIBUTIONS

L. Downie carried out all experimental work with the exception of SBF-SEM experiments (performed by N. Ferrandiz), CRISPR knock-in cell line generation (N. Ferrandiz), LBR point mutation and truncation work (M. Jones). Data analysis was done by L. Downie and S.J. Royle.

## Supplementary Information

**Figure S1.**
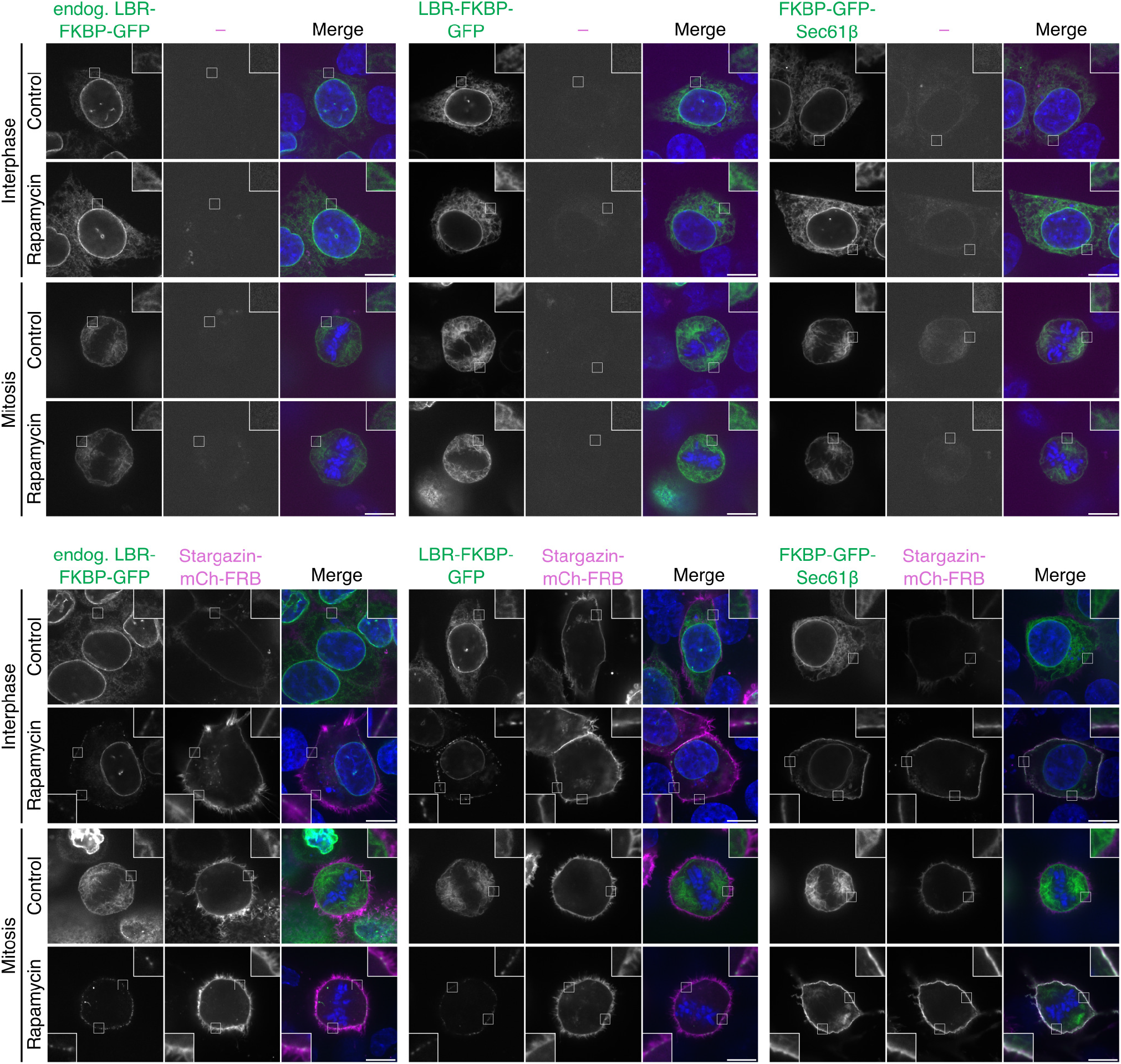
Further controls for rapamycin treatment and expression of the plasma membrane anchor construct. Example micrographs of HCT116 wild-type cells transiently expressing LBR-FKBP-GFP or FKBP-GFP-Sec61β (green) and HCT116 LBR-FKBP-GFP CRISPR knock-in cells, transiently expressing Stargazin-mCherry-FRB (magenta) as indicated and stained with DAPI (blue). Relocalized samples were treated with rapamycin (200 nM) for 30 min before fixation. Control samples were not treated with rapamycin. Scale bars, 10 μm; Insets, 3× expansion of ROI.

**Figure S2.**
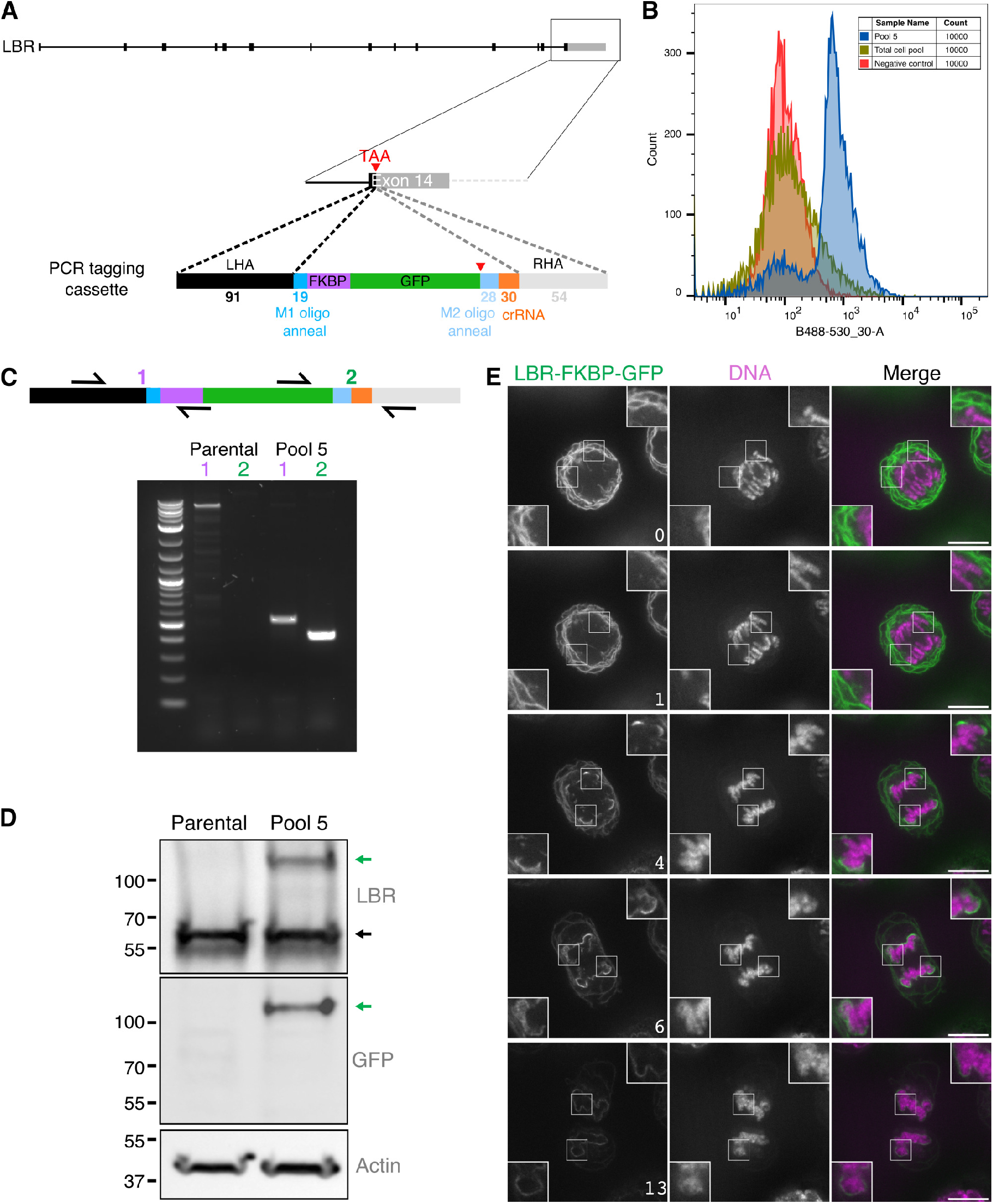
HCT116 LBR-FKBP-GFP knock-in cells. (**A**) Schematic diagram of C-terminal PCR tagging of LBR with FKBP-GFP. (**B**) FACS plots to show the collection of a mixed population of GFP positive cells. (**C**) Agarose gel to show genotyping PCR results using indicated primers. (**D**) Western blot of lysates collected from parental cells or the edited cell pool. Proteins were detected by anti-GFP or anti-LBR along with an actin loading control. Expected mass of tagged gene product (LBR-FKBP-GFP) and the untagged endogenous LBR are indicated by green and black arrows respectively. (**E**) Stills of single slices from a z-stack of live HCT116 LBR-FKBP-GFP (green) knock-in cells with SiR-DNA staining (magenta) progressing through mitosis. Time, min; scale bar, 10 μm; insets, 2× zoom.

**Figure S3.**
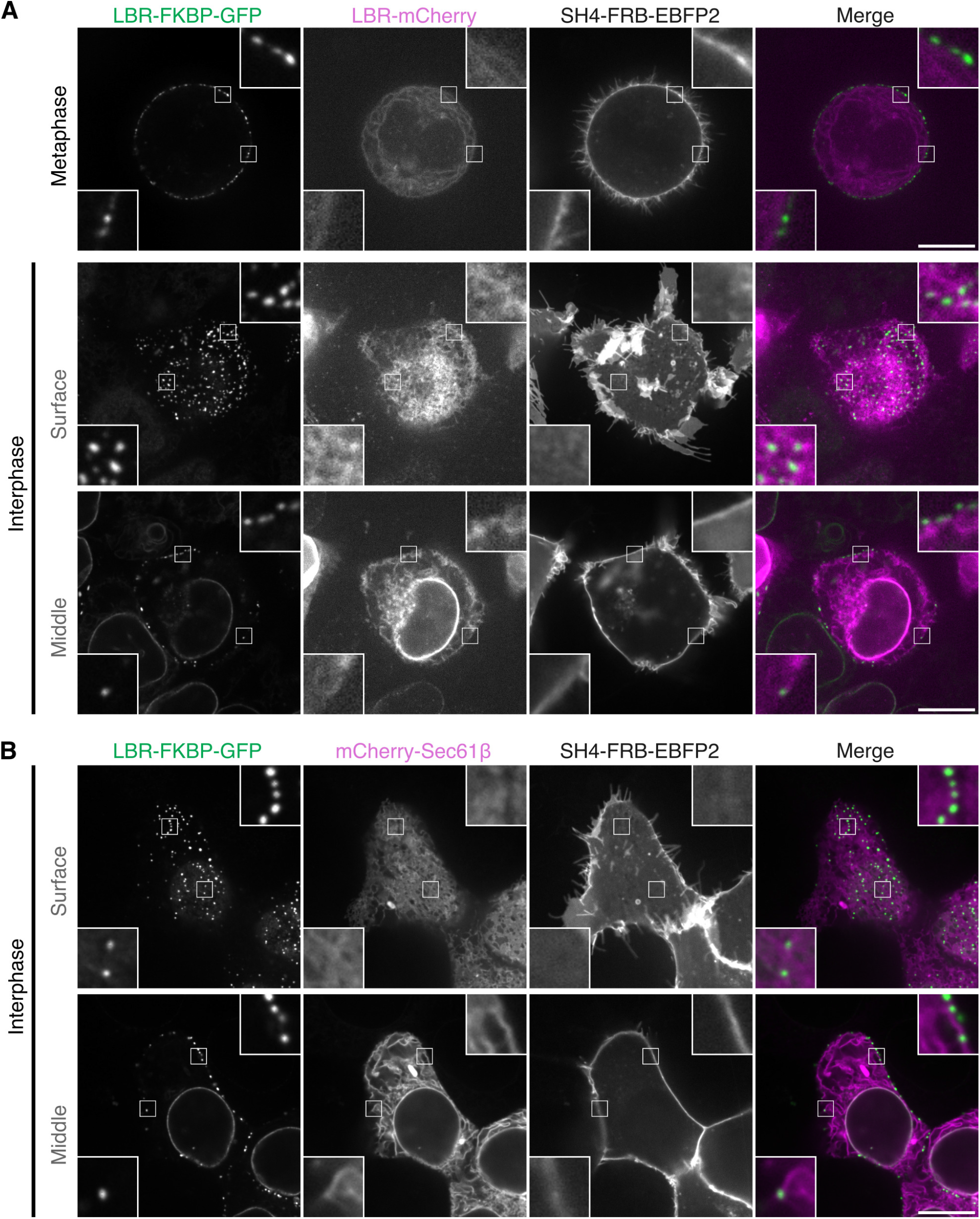
LBR-FKBP-GFP relocalization does not cluster the plasma membrane anchor or other ER proteins. Single slices from z-stacks of live HCT116 cells co-expressing LBR-FKBP-GFP (green), LBR-mCherry (**A**) or mCherry-Sec61β (**B**) (magenta) and SH4-FRB-EBFP2 (not shown in merge), treated with rapamycin (200 nM). Scale bars, 10 μm; Insets, 4× expansion of ROI.

**Figure S4.**
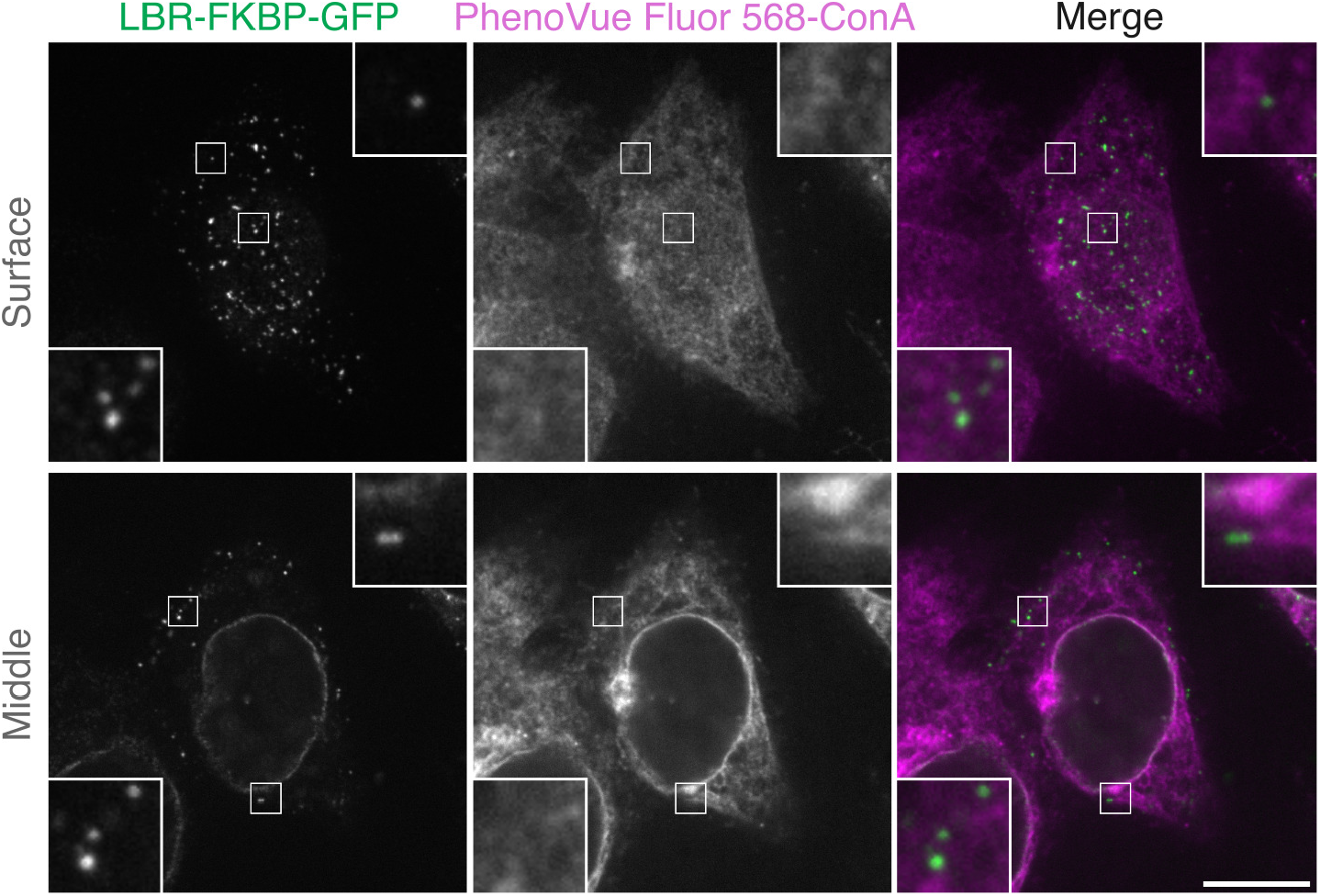
LBR-FKBP-GFP relocalization does not alter ER morphology. Single slices of fixed HCT116 cells co-expressing LBR-FKBP-GFP (green) and SH4-FRB-EBFP2 (not shown), treated with rapamycin (200 nM) for 30 min before fixation, and then stained with concanavalin A dye (PhenoVue Fluor 568-ConA, magenta). Scale bars, 10 μm; Insets, 4× expansion of ROI.

**Figure S5.**
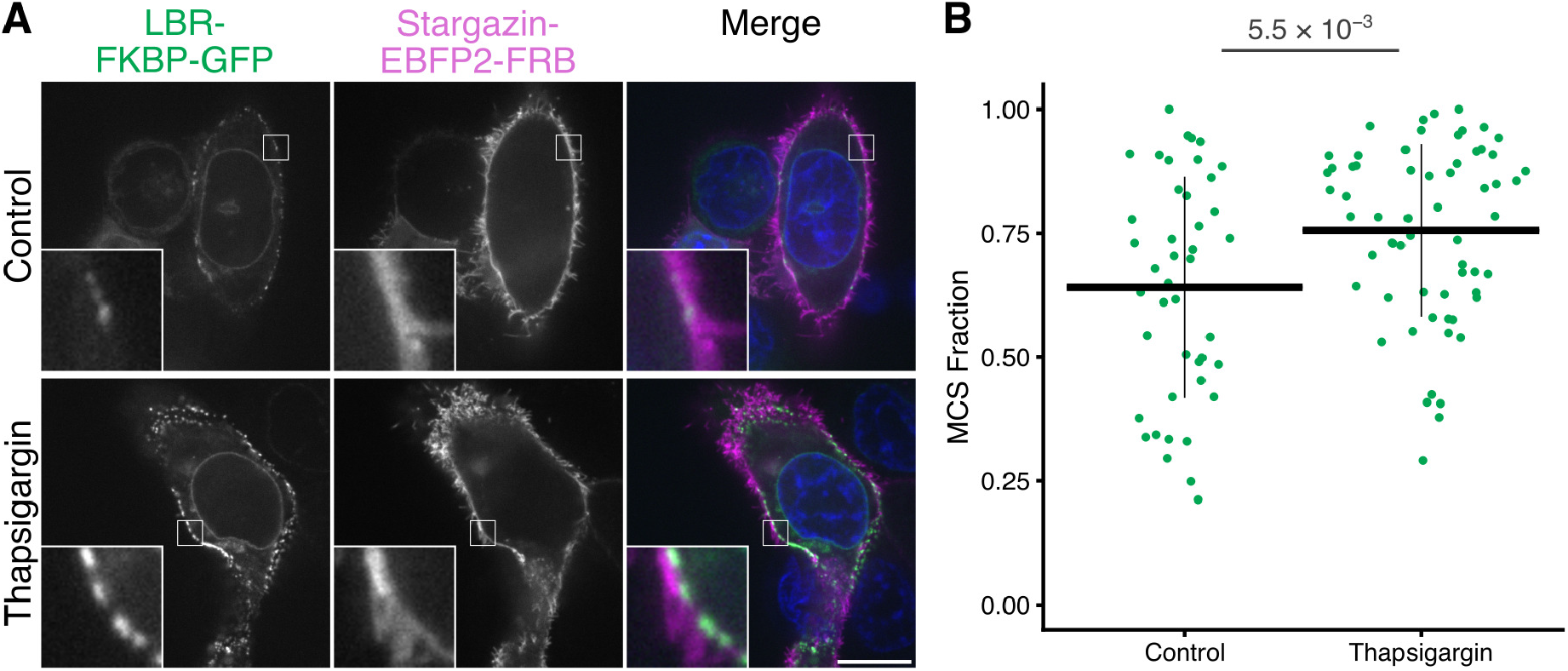
Detecting a thapsigargin-induced increase in ER-PM contact sites with relocalization of LBR-FKBP-GFP. (**A**) Typical confocal images of HCT116 cells co-expressing LBR-FKBP-GFP (green), Stargazin-EBFP2-FRB (magenta), with SiR-DNA (blue in merge) to detect DNA. Cells were treated with thapsigargin (1 μM, 20 min) or not, as indicated, before relocalization was induced with rapamycin (200 nM). Scale bar, 10 μm; Insets, 4× expansion of ROI. (**B**) Plot to show the MCS fraction of the plasma membrane profile. Spot, cells; bars, mean ± sd. P-value, Student’s t-test.

**Figure S6.**
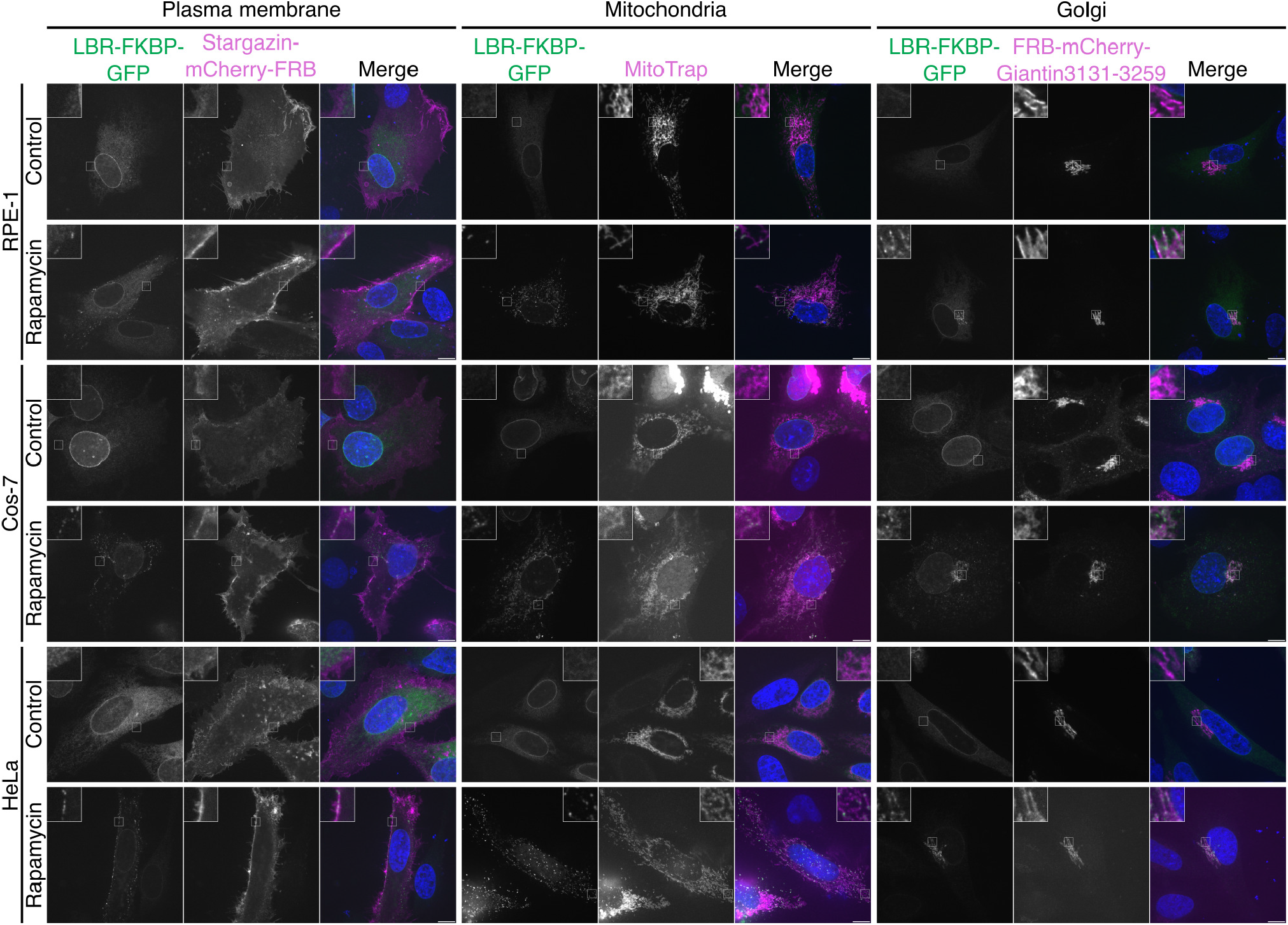
LBR-FKBP-GFP relocalizes in a similar pattern and can act as a generic marker of several ER-membrane contact types across multiple different cell types. Single slices from z-stacks of fixed RPE-1, Cos-7 and HeLa cells co-expressing LBR-FKBP-GFP (green) and Stargazin-mCherry-FRB, Mito-mCherry-FRB or FRB-mCherry-Giantin(3131-3259) (magenta), treated with rapamycin (200 nM for 30 min before fixation) where indicated and stained with DAPI (blue). Scale bars, 10 μm; Insets, 4× expansion of ROI.

**Figure S7.**
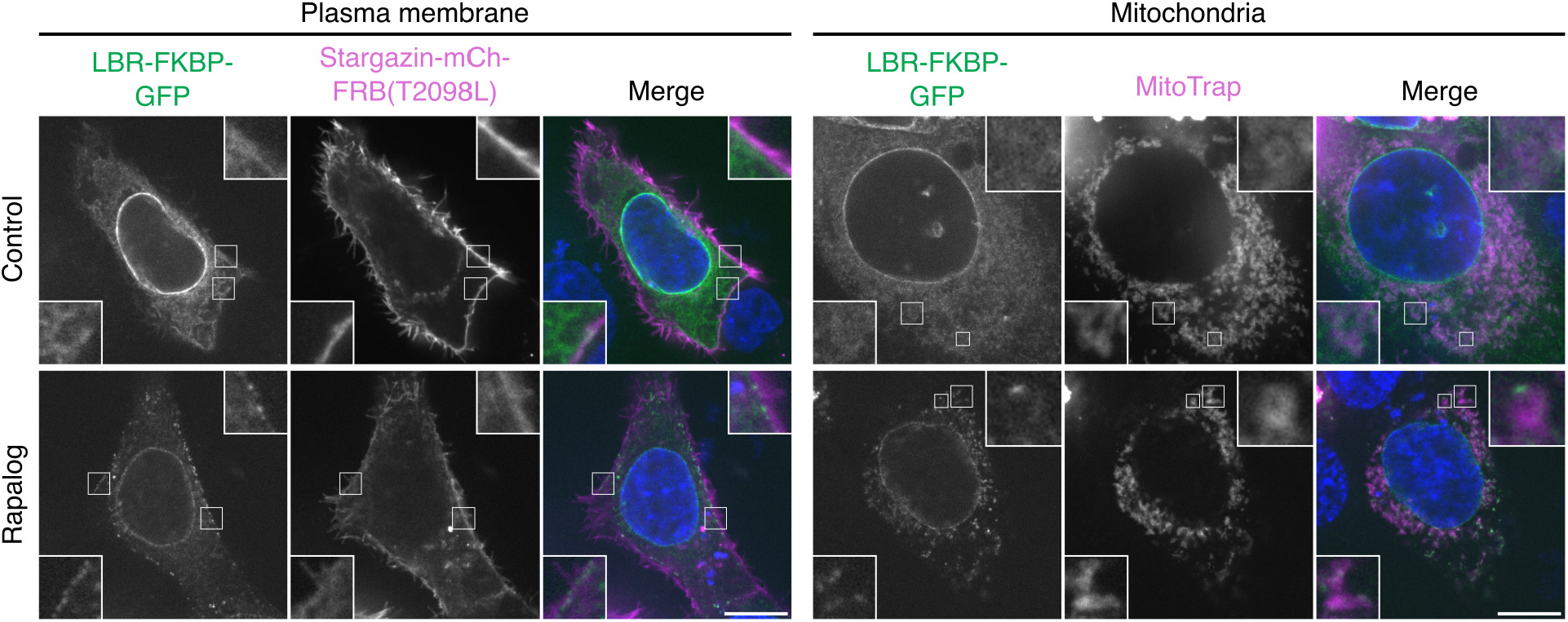
Relocalization of LBR-FKBP-GFP to ER-PM or ER-Mitochondria contact sites using rapalog. Single slices from z-stacks of fixed HCT116 cells co-expressing LBR-FKBP-GFP (green) and Stargazin-mCherry-FRB(T2098L) or MitoTrap (Mito-mCherry-FRB[T2098L]) (magenta) and stained with DAPI (blue). Where indicated, samples were treated with rapalog (5 μM) for 30 min prior to fixation. Scale bars, 10 μm; Insets, 6× expansion of smaller ROI or 3× expansion of larger ROI.

**Figure S8.**
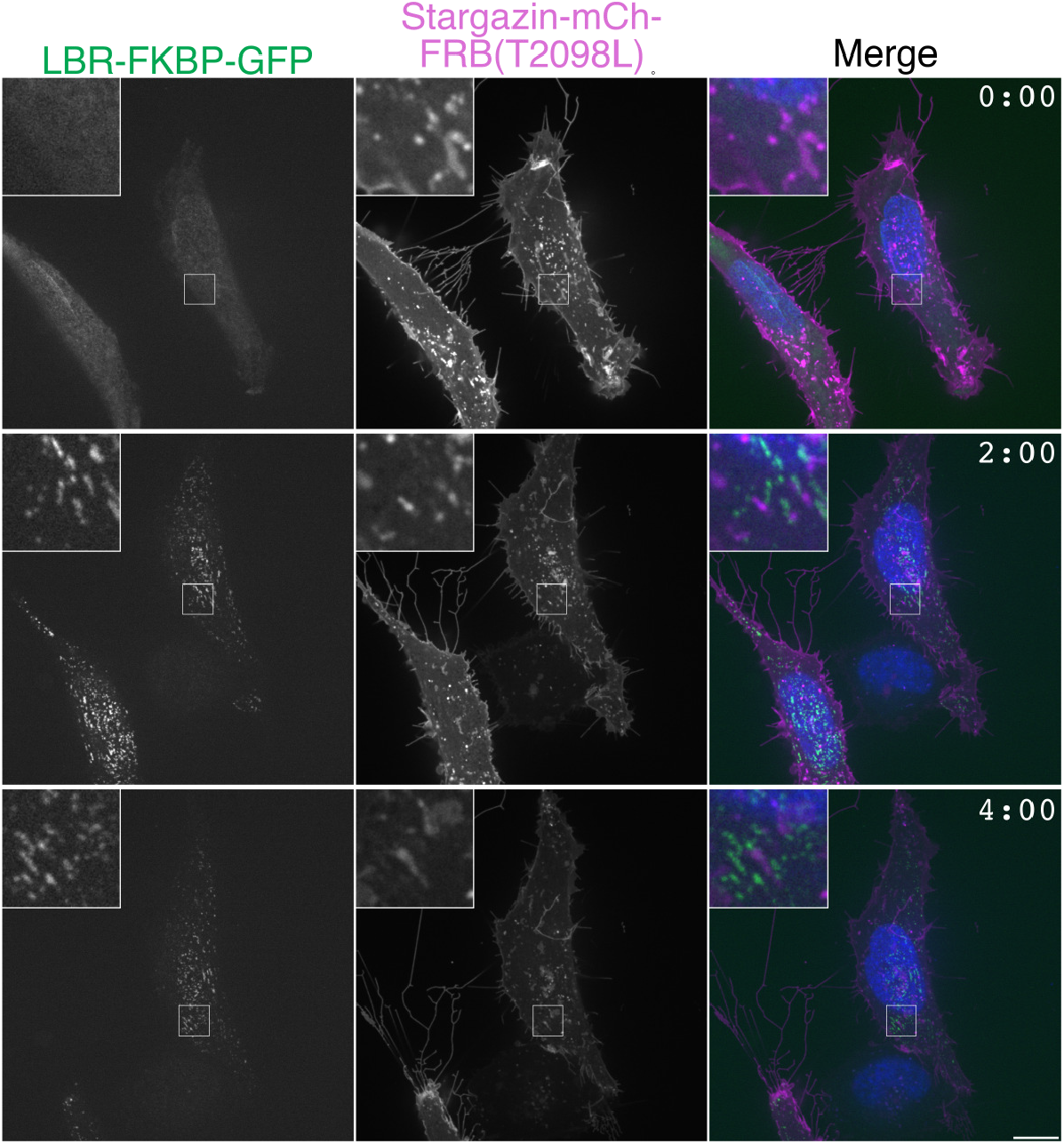
LBR-FKBP-GFP relocalization labelling long-term. Live HeLa cells transiently expressing LBR-FKBP-GFP (green) and Stargazin-mCherry-FRB(T2098L) (magenta) with SiR-DNA staining (blue). Rapalog was added to a final concentration 5 μM after capture of images at the first timepoint (0 h). Images were captured at 0, 2 and 4 h. Time is indicated in hh:mm. Scale bars, 10 μm; Insets, 4× expansion of ROI.

**Figure S9.**
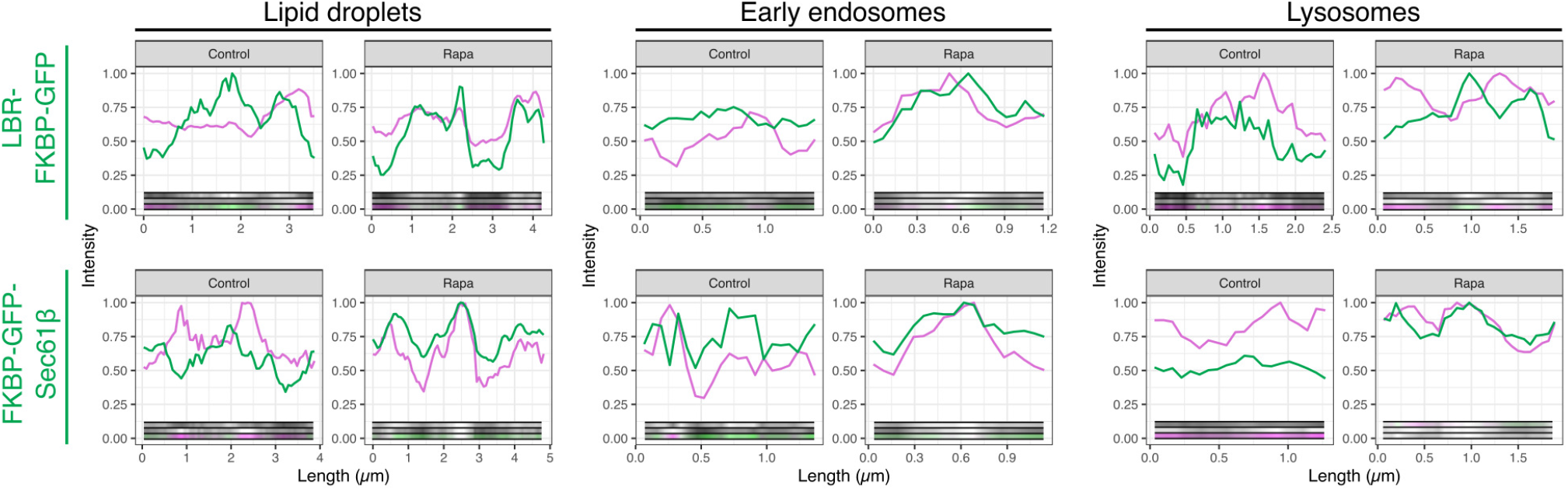
Line profiles of LBR-FKBP-GFP or FKBP-GFP-Sec61β relocalization to mark ER-lipid droplet, ER-endosome or ER-lysosome contact sites. Line profiles corresponding to the cells shown in Figure 5A. Plots show the intensity of LBR-FKBP-GFP or FKBP-GFP-Sec61bβ (green) and anchor protein (magenta) signal measured around the perimeter of the structure. Anchor proteins are: FRB-mCherry-PLIN3 (lipid droplets), FRB-mCherry-EEA1 (early endosomes), or LAMP1-mCherry-FRB (lysosomes). Insets show the line profile images.

**Figure S10.**
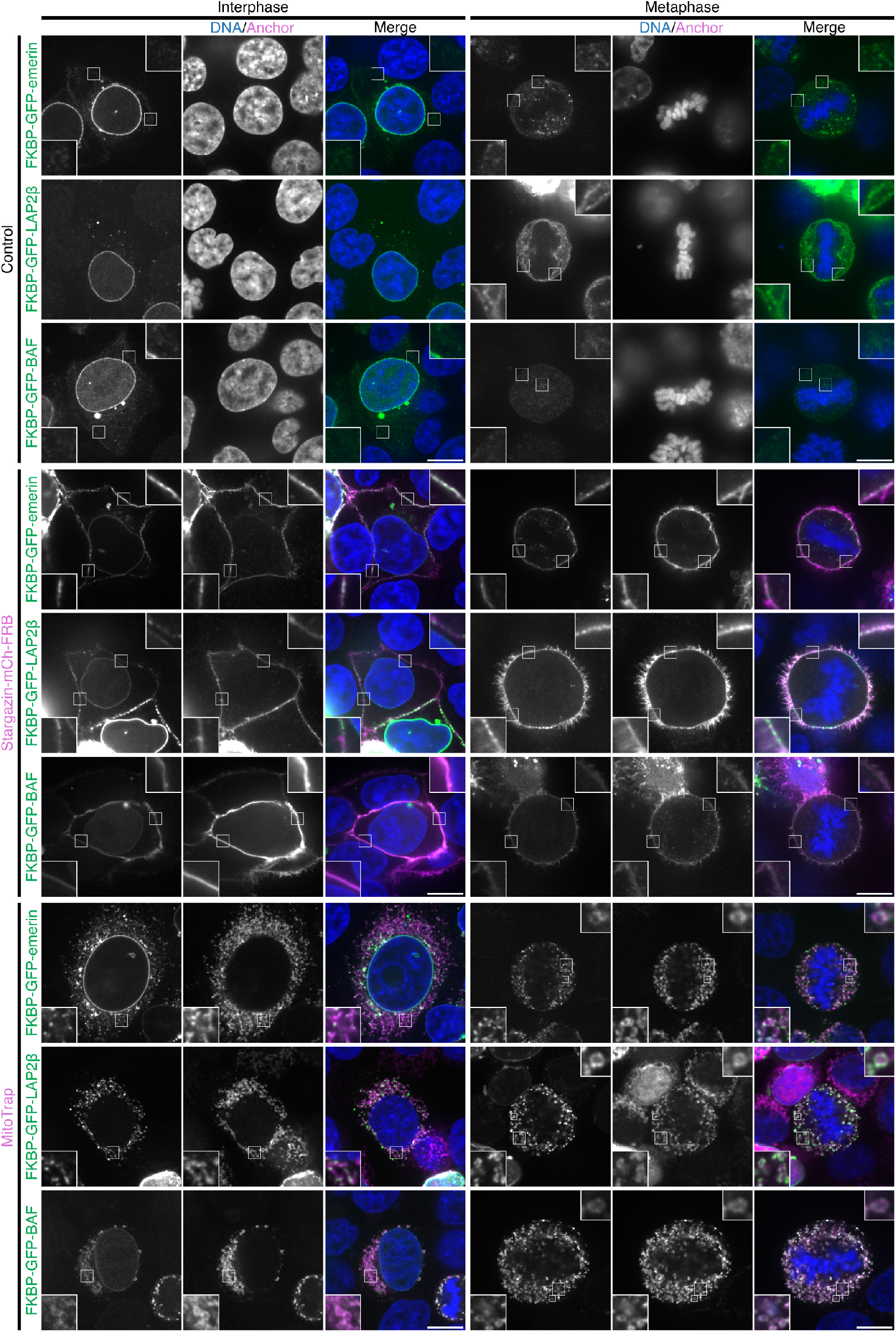
Other proteins tested do not form clusters when relocalized to the plasma membrane or mitochondria. Example micrographs of HCT116 cells co-expressing emerin, LAP2β or BAF construct tagged with FKBP-GFP at the N-terminus (green), with Stargazin-mCherry-FRB, Mito-mCherry-FRB (**C**) or a control with no FRB construct expressed (as indicated), stained with DAPI (blue). All samples were treated with rapamycin (200 nM) for 30 min prior to fixation. Shown are single slices from z-stacks of an interphase and metaphase cell. Scale bars, 10 μm; insets, 5× expansion of smaller ROI in mitochondria examples or 3× expansion of larger ROI.

**Figure S11.**
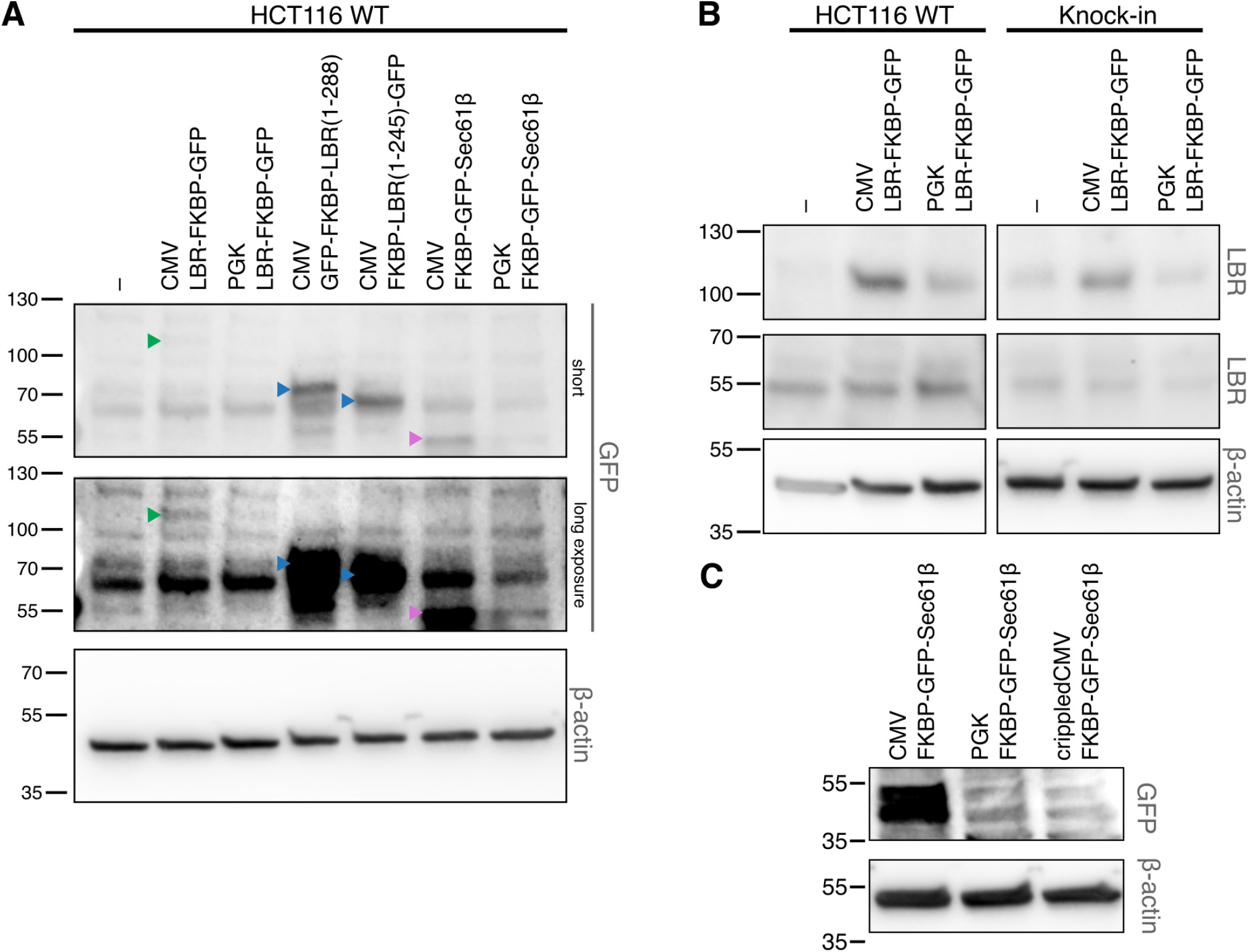
Comparing expression levels of LBR-FKBP-GFP at endogenous level and FKBP-GFP-Sec61β expressed under different promoters. (**A**) Western blot detection of lysates collected from HCT116 cells transiently expressing LBR-FKBP-GFP or FKBP-GFP-Sec61β under different promoters (CMV or PGK), LBR truncations with GFP and FKBP tags (shown in Figure 6) and a no transfection control (–). Proteins were detected by an anti-GFP antibody with β-actin loading control. Expected masses of GFP- and FKBP-tagged LBR, LBR truncations and Sec61β are indicated by green, blue and magenta arrow heads respectively. (**B**) Western blot of similarly prepared lysates collected from HCT116 or HCT116 LBR-FKBP-GFP CRISPR knock-in cells. Proteins were detected using anti-LBR antibody with β-actin loading control. The bands around the expected mass of LBR-FKBP-GFP (top) and endogenous LBR (middle) are shown. (**C**) Western blot of similarly prepared lysates from HCT116 cells transiently expressing FKBP-GFP-Sec61β under different promoters (CMV, PGK and crippledCMV) detected using an anti-GFP antibody with β-actin loading control.

**Figure S12.**
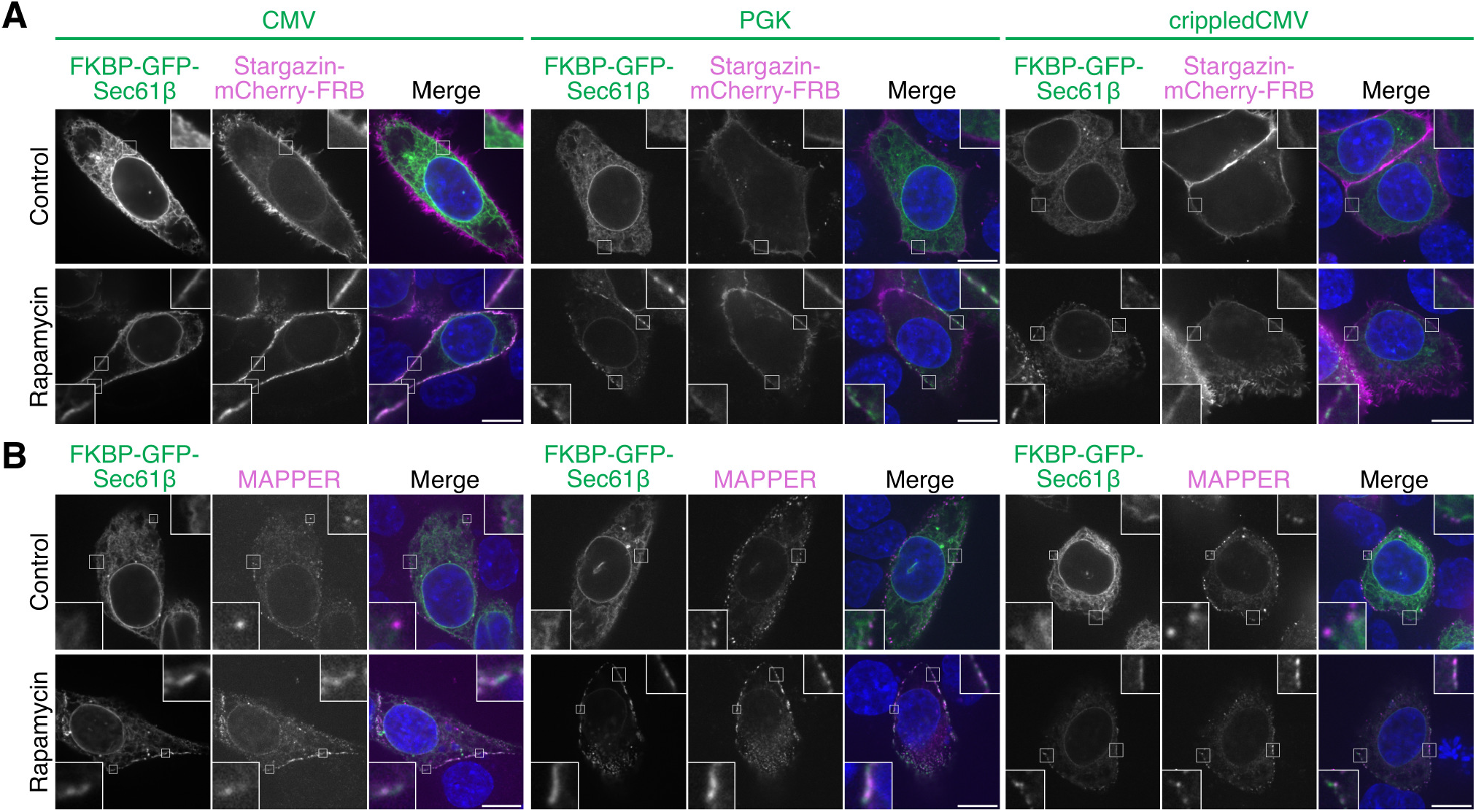
FKBP-GFP-Sec61β relocalizes to areas larger than ER-PM contact sites, even at the lowest expression levels. (**A**) Representative confocal images of relocalization of FKBP-GFP-Sec61β to Stargazin-mCherry-FRB (magenta) transiently expressed in HCT116 cells under CMV, PGK or crippledCMV promoters. (**B**) Representative confocal images of relocalization of FKBP-GFP-Sec61β to Stargazin-EBFP2-FRB transiently expressed in HCT116 cells under CMV, PGK or crippledCMV promoters. Cells additionally express mScarlet-I3-6DG5-MAPPER (magenta). Scale bars, 10 μm; insets, 6× expansion of smaller ROI or 3× expansion of larger ROI.

**Figure S13.**
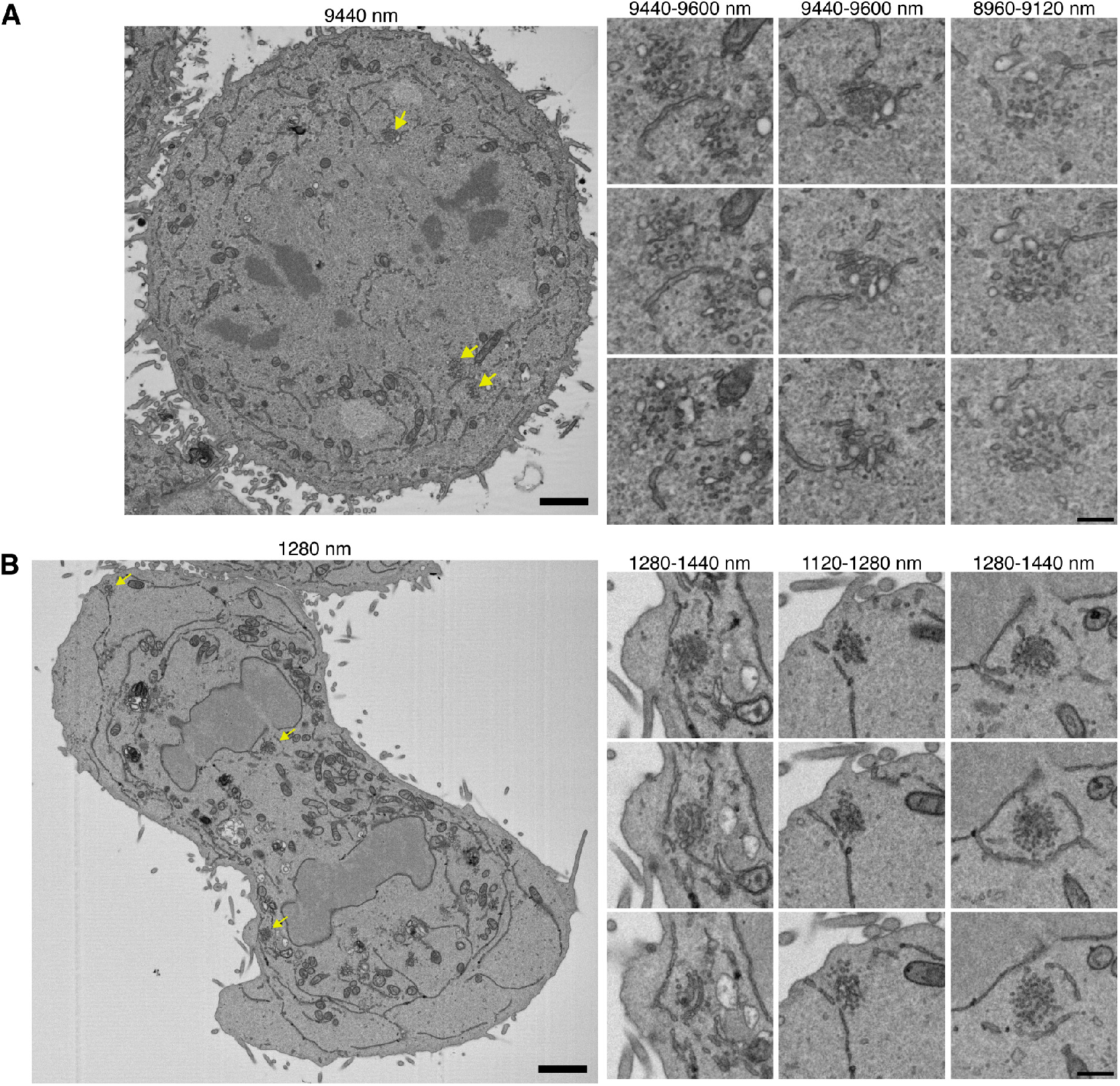
Further examples of mitotic ER-Golgi contacts by SBF-SEM. Single slices of metaphase (**A**) and telophase (**B**) HCT116 LBR-FKBP-GFP CRISPR knock-in cell SBF-SEM datasets are shown. Depth of each slice within the dataset (nm) is indicated. Example Golgi clusters are shown by yellow arrows on the full slice image. Three sequential slices of these regions (3× expansion) are shown beside. Scale bars, 2 μm and 0.5 μm on zoom region.

## Supplementary Videos

**Figure SV1.**
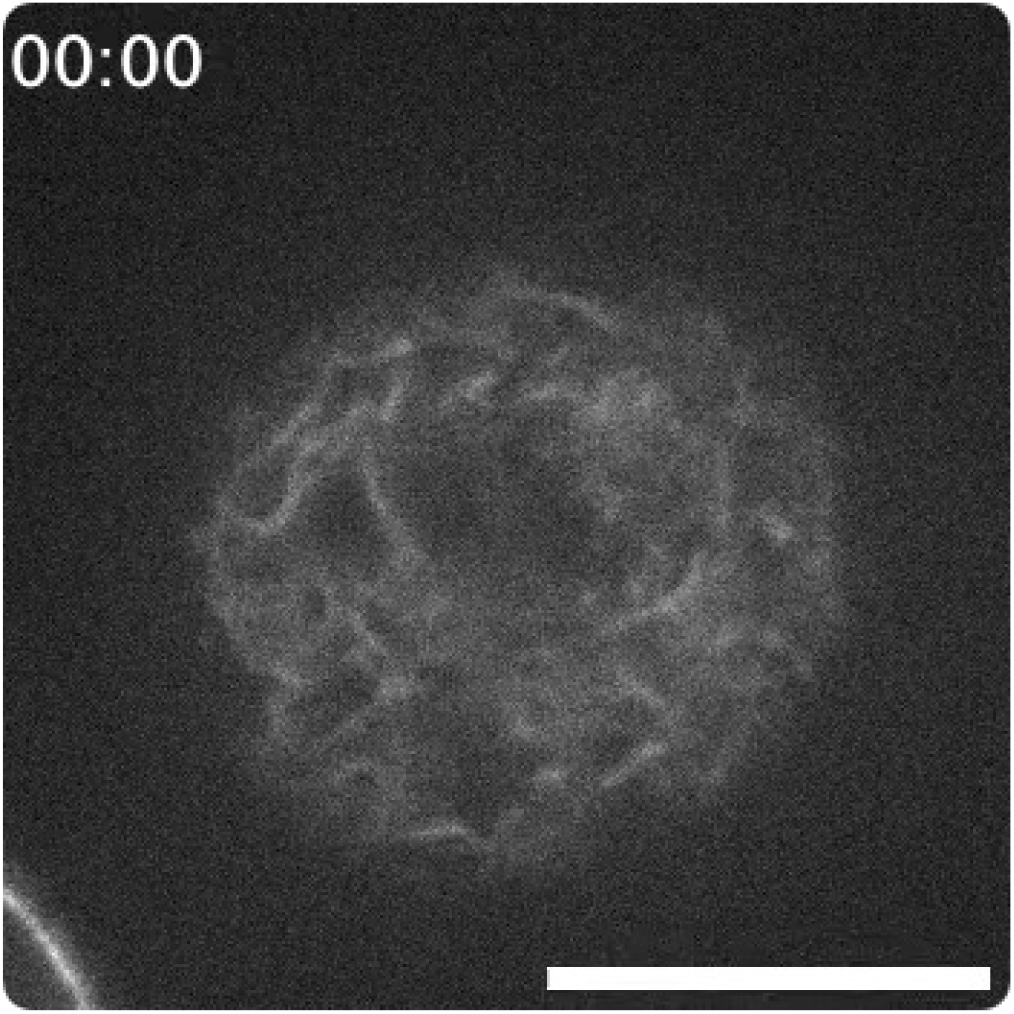
Example of induced relocalization of LBR-FKBP-GFP to the plasma membrane. Movie of a mitotic HCT116 LBR-FKBP-GFP knock-in cell co-expressing Stargazin-mCherry-FRB (not shown). Rapamycin (200 nM) is added between the first and second frame. Time, mm:ss. Playback, 10 fps. Scale bar, 10 μm.

**Figure SV2.**
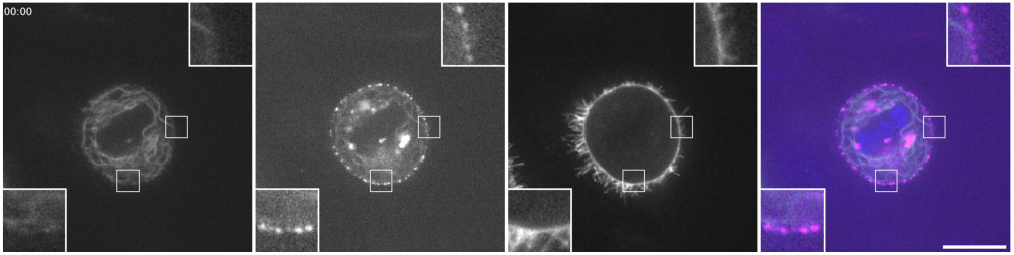
Example of LBR-FKBP-GFP relocalisation to the plasma membrane in HCT116 cells expressing MAPPER. Movie of a mitotic HCT116 cell co-expressing LBR-FKBP-GFP (green, left), mScarlet-I3-6DG5-MAPPER (magenta, second channel), Stargazin-EBFP2-FRB (third channel) and stained with SiR-DNA (blue). Rapamycin (200 nM) is added between the first and second frame. Time, mm:ss. Playback, 2 fps. Insets, 3× expansion of ROI. Scale bar, 10 μm.

**Figure SV3.**
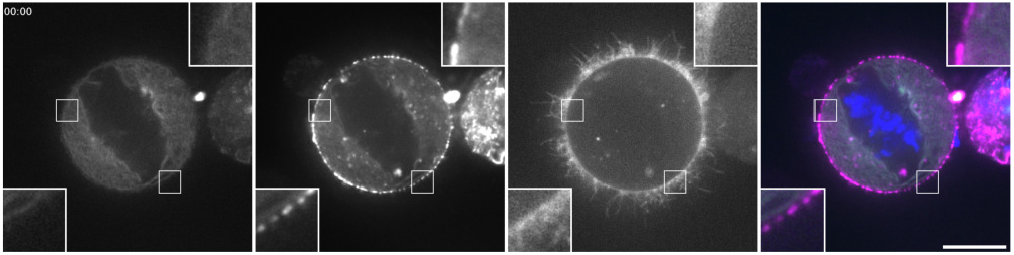
Example of LBR-FKBP-GFP relocalisation to the plasma membrane in HeLa cells expressing MAPPER. Movie of a mitotic HeLa cell co-expressing LBR-FKBP-GFP (green, left), mScarlet-I3-6DG5-MAPPER (magenta, second channel), Stargazin-EBFP2-FRB (third channel) and stained with SiR-DNA (blue). Rapamycin (200 nM) is added between the first and second frame. Time, mm:ss. Playback, 2 fps. Insets, 3× expansion of ROI. Scale bar, 10 μm.

**Figure SV4.**
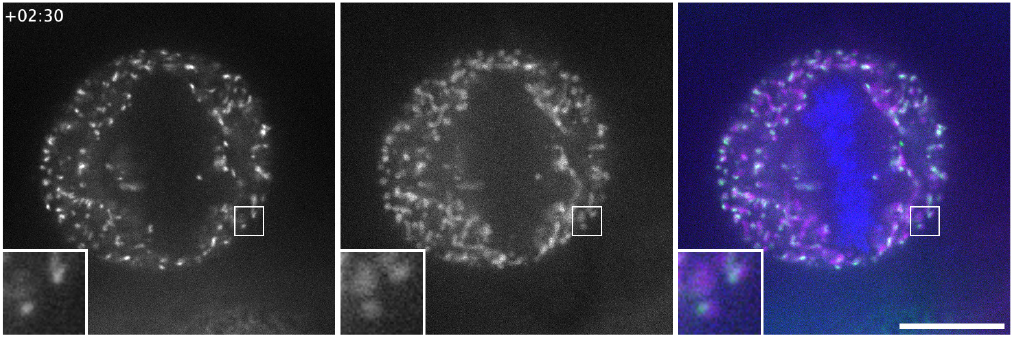
Example of induced relocalization of LBR-FKBP-GFP to mitochondria. Movie of a mitotic HCT116 LBR-FKBP-GFP (green, left) knock-in cell co-expressing MitoTrap (Mito-mCherry-FRB, magenta, middle), stained with SiR-DNA (blue). Rapamycin (200 nM) is added during frame 2. Time, mm:ss. Playback, 10 fps. Insets, 3× expansion of ROI. Scale bar, 10 μm.

**Figure SV5.**
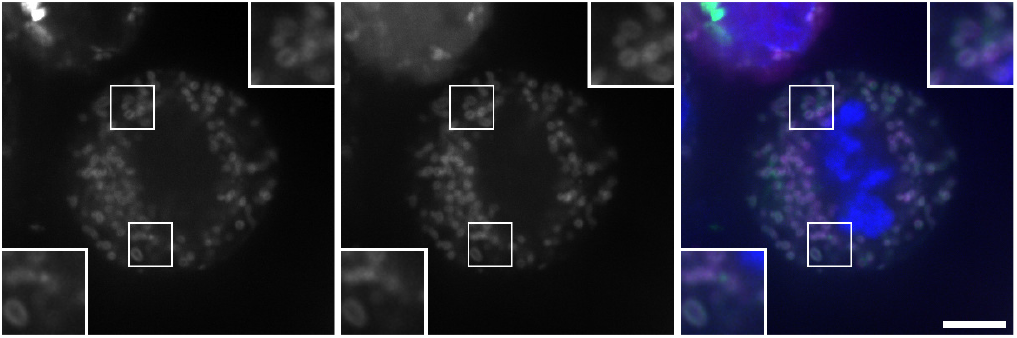
Example of induced relocalization of FKBP-GFP-Sec61β to mitochondria. Movie of a mitotic HCT116 cell co-expressing FKBP-GFP-Sec61β (green, left), MitoTrap (Mito-mCherry-FRB, magenta, middle), stained with SiR-DNA (blue). Rapamycin (200 nM) is added between the first and second frame. Time, mm:ss. Playback, 1 fps. Insets, 2× expansion of ROI. Scale bar, 5 μm.

**Figure SV6.**
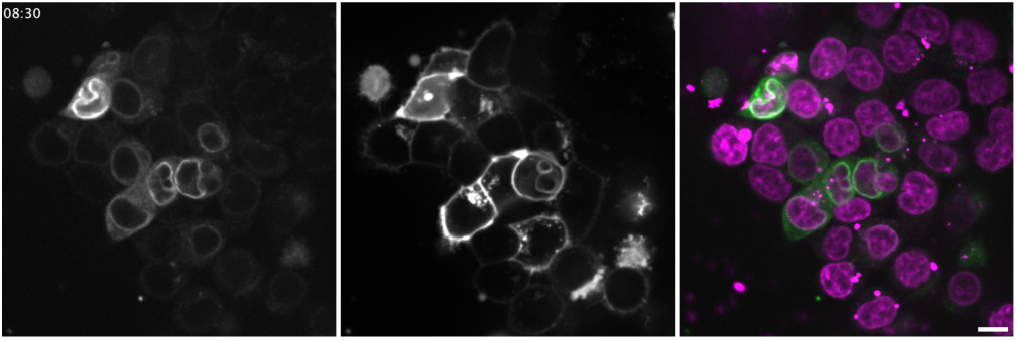
LBR-FKBP-GFP imaging long-term in live HCT116 cells. Live HCT116 cells transiently expressing LBR-FKBP-GFP (green, left) and Stargazin-mCherry-FRB(T2098L) (middle) with SiR-DNA staining (magenta). Videos were captured in a single z-slice at 30 min intervals. Time is indicated in hh:mm. Playback, 2 fps. Scale bar, 10 μm.

**Figure SV7.**
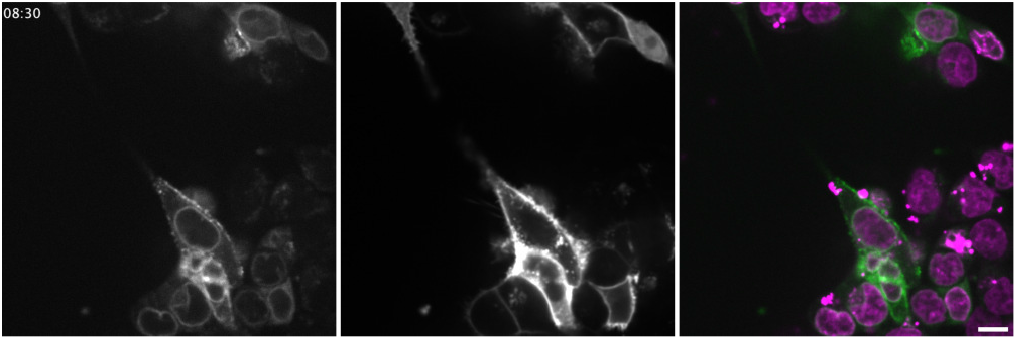
LBR-FKBP-GFP relocalisation long-term in live HCT116 cells. Live HCT116 cells transiently expressing LBR-FKBP-GFP (green, left) and Stargazin-mCherry-FRB(T2098L) (middle) with SiR-DNA staining (magenta). Rapalog was added to a final concentration of 5 μM between the first and second frame. Videos were captured in a single z-slice at 30 min intervals. Time is indicated in hh:mm. Playback, 2 fps. Scale bar, 10 μm.

**Figure SV8.**
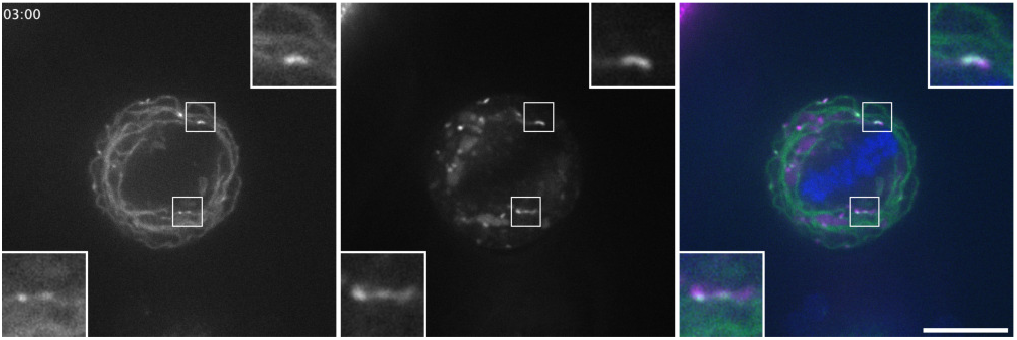
Golgi HCT116 video. Movie of a mitotic HCT116 cell co-expressing LBR-FKBP-GFP (green, left), FRB-mCherry-Giantin3131-3259 (magenta, middle) and stained with SiR-DNA (blue). Rapamycin (200 nM) is added between the first and second frame. Time, mm:ss. Playback, 2 fps. Insets, 3× expansion of ROI. Scale bar, 10 μm.

